# A molecularly defined NAcSh D1 subtype controls feeding and energy homeostasis

**DOI:** 10.1101/2023.02.27.530275

**Authors:** Yiqiong Liu, Ying Wang, Zheng-dong Zhao, Guoguang Xie, Chao Zhang, Renchao Chen, Yi Zhang

## Abstract

Orchestrating complex behavioral states, such as approach and consumption of food, is critical for survival. In addition to hypothalamus neuronal circuits, the nucleus accumbens (NAc) also plays an important role in controlling appetite and satiety in responses to changing external stimuli. However, the specific neuronal subtypes of NAc involved as well as how the humoral and neuronal signals coordinate to regulate feeding remain incompletely understood. Here, we deciphered the spatial diversity of neuron subtypes of the NAc shell (NAcSh) and defined a dopamine receptor D1(Drd1)- and *Serpinb2-*expressing subtype located in NAcSh encoding food consumption. Chemogenetics- and optogenetics-mediated regulation of *Serpinb2*^+^ neurons bidirectionally regulates food seeking and consumption specifically. Circuitry stimulation revealed the NAcSh^Serpinb2^→LH^LepR^ projection controls refeeding and can overcome leptin-mediated feeding suppression. Furthermore, NAcSh *Serpinb2*^+^ neuron ablation reduces food intake and upregulates energy expenditure resulting in body weight loss. Together, our study reveals a neural circuit consisted of molecularly distinct neuronal subtype that bidirectionally regulates energy homeostasis, which can serve as a potential therapeutic target for eating disorders.

## Main

Feeding is a complicated motivational and emotional behavior required for survival and is known for its extraordinary ability to adapt in response to environmental changes^1, 2^. The obesity epidemic coupled with the increased population with eating disorders, such as binge eating and anorexia, underscores the urgent need for understanding the mechanisms controlling feeding behavior^3, 4^.

Energy homeostasis is a complicated process largely controlled by neuronal circuits in the hypothalamus and brainstem^5–7^, while reward and motivation of food intake are mainly processed in the limbic regions^8^ and cerebral cortex^9, 10^. The hypothalamus, with highly heterogenous neuronal composition^11^, plays a critical role in controlling feeding behavior^12, 13^. The core of the hypothalamic control system in the arcuate nucleus (Arc) are comprised of two neuronal populations, the AgRP neurons and the Pro-opiomelanocortin (POMC) neurons. These two types of neurons exert almost opposite functions in regulating feeding, with feeding related hormones such as ghrelin and leptin, coordinately mediate the sensations of appetite and satiety leading to behavioral response^14, 15^. Leptin performs anorexic function by acting on leptin receptors (LepR) in the hypothalamus^16, 17, 18^, In addition to the Arc, LepR is also highly expressed in lateral hypothalamus (LH)^19^, which receives neural inputs from Arc and dorsomedial (DM) nucleus making it a key brain region for feeding behavior regulation. However, the specific inputs outside of hypothalamus regulating LH^LepR^ neurons in the context of feeding is not known.

Food is naturally rewarding and typically acts on the reward pathways in the brain. The nucleus accumbens (NAc) is a key component of the basal ganglion circuitry, which integrates information from cortical and limbic regions to direct feeding behaviors^20^.It has been shown that pleasure of food consumption is connected with the rostral dorsomedial part of the NAc shell (NAcSh)^21^, while the motivation to eat or incentive salience is connected with both shell and core regions^22^. In recent years, several studies have analyzed the role of the mediodorsal NAcSh in feeding^20, 23, 24^ and revealed that activation of the dopamine receptor 1 expressing medium spiny neurons (D1-MSNs)^25, 26^ projecting to the lateral hypothalamus (LH)^21, 24^ or ventral tegmental area (VTA) ^27^ stops ongoing food consumption. However, other studies showed that D1-MSNs activity was enhanced during appetitive phase^28^ as well as consumption^29^. Although temporally distinct phases of feeding behavior, such as food seeking, food evaluation and consumption, could potentially account for such discrepancy, the heterogeneity of NAc neurons^30^ could underlie the discrepancies considering that different studies might have manipulated different neuron subtypes with opposing functions.

With the application of single cell RNA-seq and spatial transcriptome techniques, it is now possible to decipher the neuron heterogeneity of different brain regions^31–35^. As a result, different molecularly and spatially distinctive neuron subtypes have been identified, making the study of neuronal circuit and subtype-specific functions of specific behaviors possible.

To identify NAc neuron subtypes that control feeding behavior, we analyzed multiplexed error-robust fluorescence in situ hybridization (MERFISH) dataset of NAcSh and identified that the *Serpinb2*-expressing D1-MSN, one of the four D1-MSNs subtypes located in NAcSh, dictates food seeking, goal-directed motivation, and food consumption. In addition, through viral tracing and terminal stimulation, we found that the NAcSh^Serpinb2^→LH^LepR^ neuronal circuit modulates feeding behavior and overcomes leptin-mediated feeding suppression. Furthermore, we showed that ablation of the *Serpinb2^+^*neurons decreases food consumption and increases energy expenditure, resulting in bodyweight loss in mice. Taken together, our study reveals a molecularly distinct accumbal-to-lateral hypothalamic neural circuit whose activity can modulate leptin-mediated effects in LH leading to the control of food consumption and bodyweight gain.

## Result

### *Serpinb2*^+^ neurons are activated in refeeding process

The NAc is a critical component of the basal ganglion circuitry, which receives and integrates information from cortical and limbic regions for actions. Previous studies have linked feeding behavior to the NAcSh^20, 21, 24, 27, 28, 36^. D1-MSNs, but not D2-MSNs, provide the dominant source of accumbal inhibition to the LH and regulate complex feeding behaviors through LH GABA neurons^20, 28^. Giving the controversial findings of D1-MSNs in feeding^20, 29, 37^, it is critical to examine neuron subtype effect of NAcSh as the discrepancy could be caused by manipulation of different neuronal subtypes of NAcSh in the different studies.

To identify the D1 neuronal subtypes located in NAcSh, we integrated the single cell and spatial transcriptomic data of NAc^30^ by analyzing 253 selected genes and the coronal sections between bregma 1.94 mm to 0.74 mm to cover the entire NAcSh region. Clustering analysis of the MERFISH data identified four major cell populations representing D1-MSNs named as D1_1 to D1_4 (Fig. 1a), which are respectively marked by *Stard5*, *Tac2*, *Spon1* and *Serpinb2* (Fig. 1b). Single molecule FISH (smFISH) further confirmed that these neuronal subtypes are mainly located in the medial dorsal NAcSh, and are predominantly *Drd1*^+^ neurons (Fig. 1c, d, Extended Data Fig.1 a-c).

**Fig. 1.**
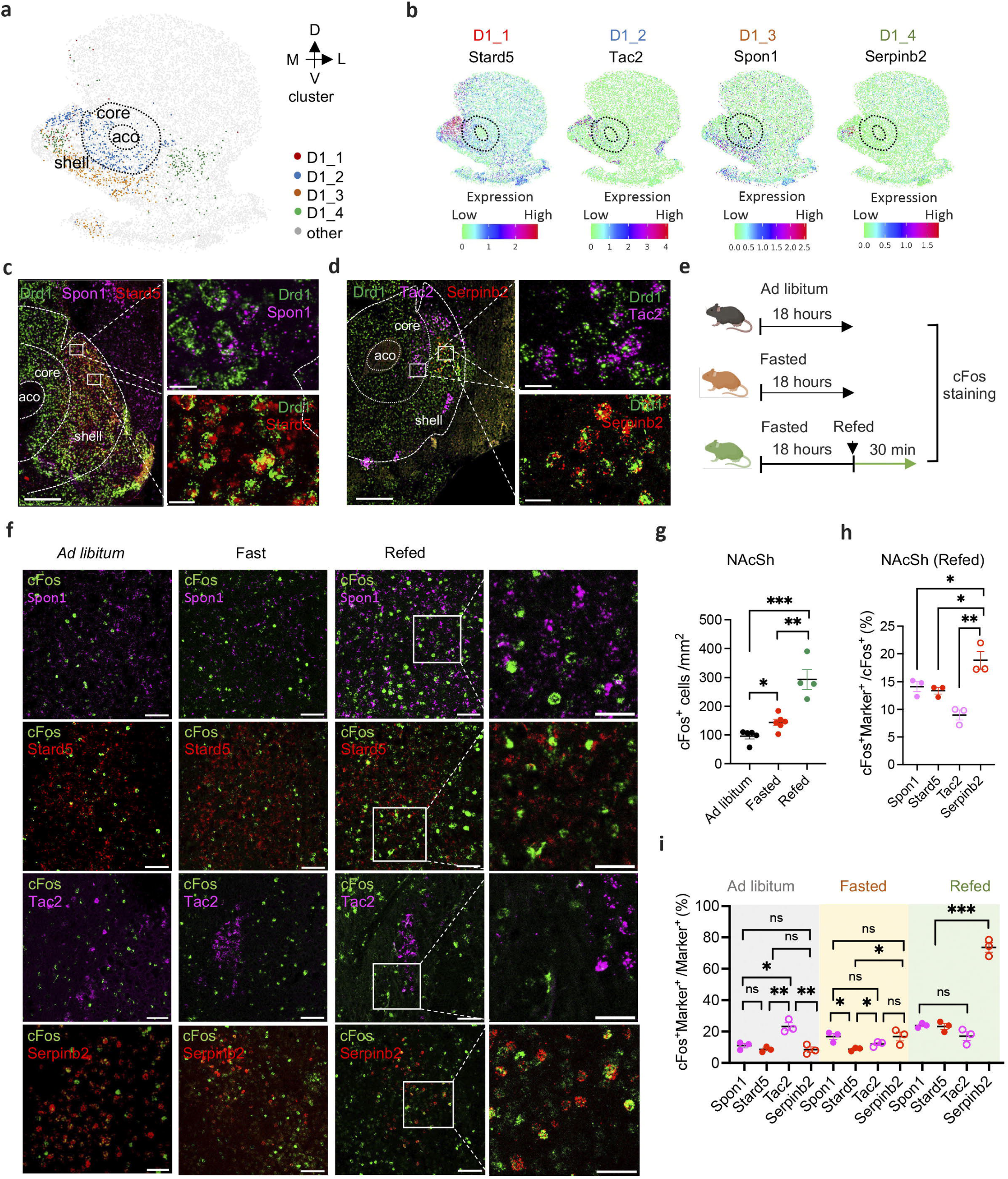
NAcSh *Serpinb2*^+^ neurons are activated in refeeding process *in vivo*. **a,** Spatial patterns of D1 MSN subtypes in NAcSh. Dotted lines circle anterior commissure olfactory limb (aco) and NAc core. The dorsal-ventral (DV) and medial-lateral (ML) axes are indicated. **b,** Spatial expression patterns from MERFISH dataset of *Stard5*, *Tac2*, *Spon1*and *Serpinb2.* These genes are able to specifically label distinct neuronal subtypes in medial dorsal NAc shell. Expression level is color-coded. **c,** RNA *in situ* hybridization showing *Spon1* and *Stard5* expression in the medial part of the NAcSh. Boxed regions in left panels are enlarged and shown in the right panels. Scale bar: 500 μm (left), 20 μm (right). **d,** RNA *in situ* hybridization showing *Tac2* and *Seprinb2* expression in the medial part of the NAcSh. Boxed regions in left panels are enlarged and shown in the right panels. Scale bar: 500 μm (left), 20 μm (right). **e,** Schematic representation of the experimental design for the *ad libitum*, fast and refeeding groups. **f,** Co-expression of *Spon1, Stard5*, *Tac2* and *Serpinb2* mRNA with *cFos* mRNA in NAc of *ad libitum*, fast and refeeding states. Representative images showing the co-localization of *cFos* (green), *Spon1* (magenta), *Stard5* (red), *Tac2* (magenta) and *Serpinb2* (red) expressing neurons. Scale bar: 50 μm. **g,** The average number of *cFos^+^* neurons in the dorsal medial NAcSh at *ad libitum*, fast and refeeding states. (n = 5 sections from three mice of each group, one-way ANOVA, with Tukey’s multiple comparisons). **h,** The percentages of activated *Spon1, Stard5*, *Tac2* or *Serpinb2* -expressing neurons in the total *cFos^+^* neurons in dorsal medial NAcSh in refeeding state. (n = 5 sections from three mice of each group, one-way ANOVA, with Tukey’s multiple comparisons). **i,** The percentages of activated *Spon1, Stard5*, *Tac2* or *Serpinb2* -expressing neurons in the total *Spon1, Stard5*, *Tac2* or *Serpinb2*-expressing neurons at *ad libitum*, fast and refeeding states. (n = 5 sections from three mice of each group, one-way ANOVA, with Tukey’s multiple comparisons). All error bars represent mean ± SEM. ***P ≤ 0.001, **P ≤ 0.01, *P ≤ 0.05; ns, P > 0.05.

To determine whether any of these D1-MSNs subtypes are involved in mediating feeding behavior, we asked whether they are activated in response to feeding by monitoring the *cFos* expression under three conditions: *ad libitum* access to food, after 18 hours of fast, and refeeding (Fig. 1e). By counting *cFos*^+^ neurons that co-express *Stard5*, *Tac2*, *Spon1* or *Serpinb2* in the medial dorsal NAcSh of smFISH (Fig. 1f), we found that the D1-MSNs subtypes responded differently to the conditions. While both fast and refeeding conditions generally increased neuronal activity compared to the *Ad libitum* condition as indicated by the increased *cFos*^+^ neuron numbers (Fig. 1g), the *Serpinb2*^+^ neurons exhibited the highest percentage of activation during refeeding with *Serpinb2*^+^ neurons accounting for 18% of total *cFos*^+^ neurons and about 70% of *Serpinb2*^+^ neurons are *cFos*^+^ (Fig. 1h, i). In contrast, only 20% of *Stard5^+^*, *Tac2^+^* and *Spon1^+^* neurons were activated in the refeeding condition (Fig. 1i). Collectively, these data indicate that the *Serpinb2*^+^ neurons are the main D1-MSN subtype in NAcSh activated in response to the refeeding process.

### The *Serpinb2*^+^ neurons respond to eating behavior

To study the role of *Serpinb2*^+^ neurons in feeding, we generated a *Serpinb2*-Cre mouse line (Extended Data Fig. 2a, b). We validated this mouse model by crossing with the Ai9 mouse line, which expresses tdTomato (tdT) fluorescence following Cre-mediated recombination, and observed about 90% colocalization of tdT signal with endogenous *Serpinb2* mRNA signal, which is consistent with endogenous *Serpinb2* expression pattern in the NAcSh as shown by the Allen Brain Atlas RNA *in situ* data (Extended Data Fig. 2c). Thus, our *Serpinb2*-Cre mouse line is suitable for studying *Serpinb2*^+^ neurons.

To determine whether *Serpinb2^+^* neurons are indeed involved in regulating feeding behavior, we used fiber photometry to monitor *Serpinb2*^+^ neuronal activity of freely moving mice during food seeking and consumption. To this end, a Cre-dependent AAV expressing the calcium reporter GCaMP7s was delivered to the NAcSh of the *Serpinb2*-Cre mice by stereotaxic injection (Fig. 2a), while AAV expressing eYFP was used as a negative control (Extended Data Fig. 3a1), followed by the implantation of an optic cannula above the NAcSh (Fig. 2b). In parallel, we implanted cannula to the *Tac2*-Cre and *Drd1*-Cre mice to monitor the NAcSh *Tac2^+^* neurons and the total NAcSh *Drd1^+^* neurons activities during feeding (Extended Data Fig. 3b1, Fig. 2b). Three weeks after the surgeries, we performed fluorescence recordings during the feeding process in home cage (Fig. 2c). Consistent with the *cFos* analysis, the activity of the *Serpinb2*^+^ neurons increased immediately when eating begins, and gradually declined when eating finishes, which is in contrast to the sniffing of an object (Fig. 2d1-d3, e1-e3). To quantify the *Serpinb2*^+^ neuronal activity at different events, we averaged the calcium signal of each animal and observed that the signals are significantly increased after the animal starting to eat chow (Fig. 2d4, left) but not the object sniffing (Fig. 2d4, right). While the signals are significantly decreased after the animal ending eating (Fig. 2e4, left) but not when leaving the object (Fig. 2e4, right). These results indicate that *Serpinb2*^+^ neuronal activity increases when starting eating the chow and decreases when ending eating. In contrast, we did not capture any obvious activity changes of the *Tac2*-Cre mice during feeding or interacting with the object (Extended Data Fig. 3b1-b3), indicating that the *Tac2^+^*neuronal subtype does not respond to feeding.

**Fig. 2.**
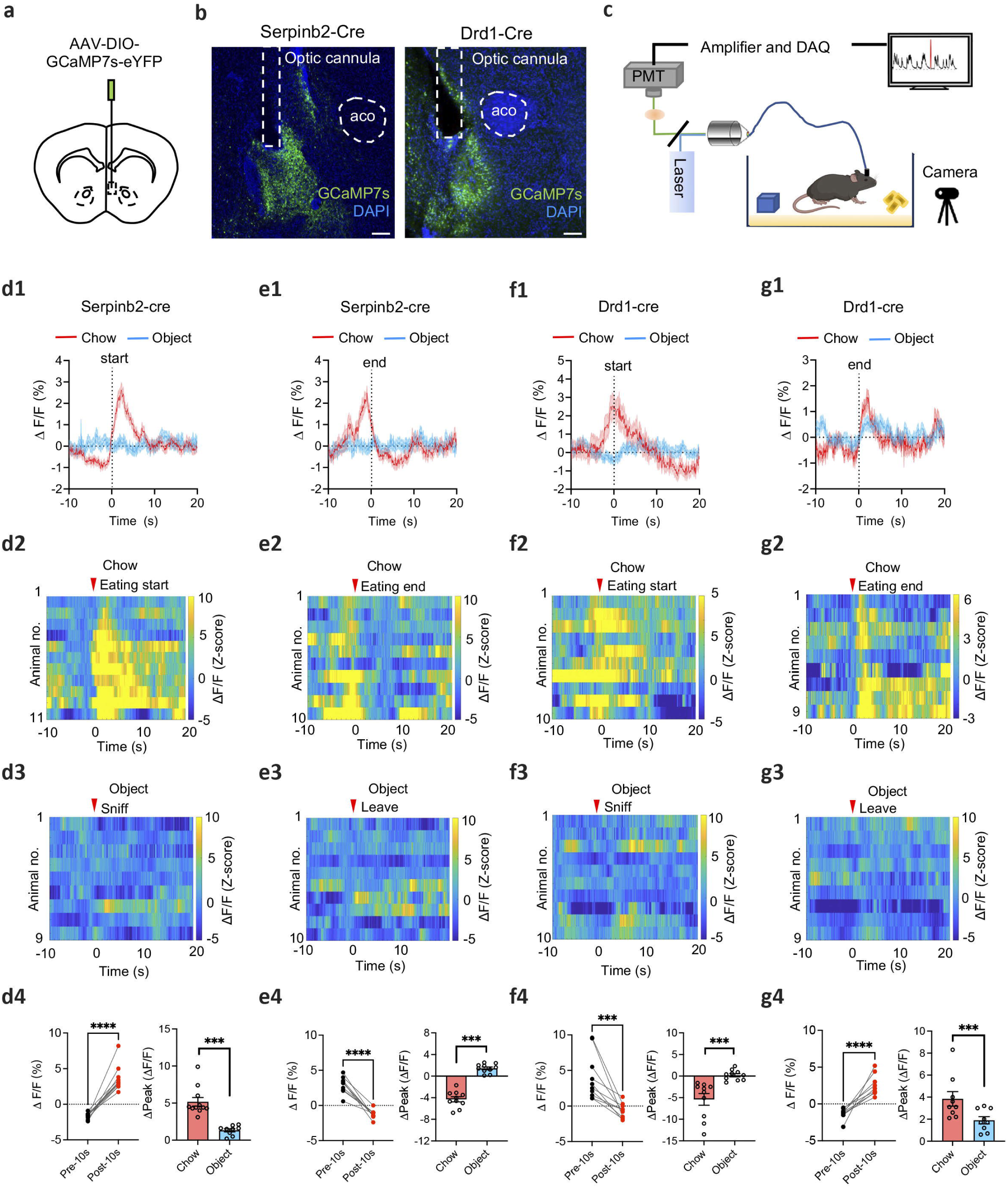
The activity of *Serpinb2*^+^ neurons is increased during feeding. **a,** Diagram of the injection site of AAV-DIO-GCaMP7s virus in NAcSh. **b,** Validation of GcaMP7s expression and implantation of optic cannula in *Serpinb2*-Cre mice (left) and *Drd1*-Cre mice (right). Scale bar: 200 μm. **c,** A schematic setup for recording form NAcSh GCaMP-expressing neurons in a freely behaving mouse. **d1-d4,** Average peri-stimulus histograms of Ca^2+^ signals of *Serpinb2*^+^ neurons. Feeding bouts, <10s; dash line, eating start point (**d1**). Corresponding heat maps of ΔF/F (Z-score) Ca^2+^ responses per animal before and after the onset of eating chow (**d2**) and sniffing object (**d3**). Averaged ΔF/F (%) is represented in the dot plot (**d4, left**). Averaged Δpeak ΔF/F is represented in the bar graph (**d4, right**). n=11 mice. **e1-e4,** Average peri-stimulus histograms of Ca^2+^ signals of *Serpinb2*^+^ neurons. Dash line, eating end point (**e1**). Corresponding heat maps of ΔF/F (Z-score) Ca^2+^ responses per animal before and after the offset of end eating chow (**e2**) and leaving object (**e4**). Averaged ΔF/F (%) is represented in the dot plot (**e4, left**). Averaged Δpeak ΔF/F is represented in the bar graph (**e4, right**). n=10 mice. **f1-f4,** similar to **d1-d4** but are recorded for the Drd1-Cre mice. **g1-g4,** similar as **e1-e4** but are recorded for the Drd1-Cre mice. ****P ≤ 0.0001, ***p<0.001; **, p<0.01; *, p<0.05; ns, p>0.05, unpaired t-test. Data are represented as mean ± SEM.

**Fig. 3.**
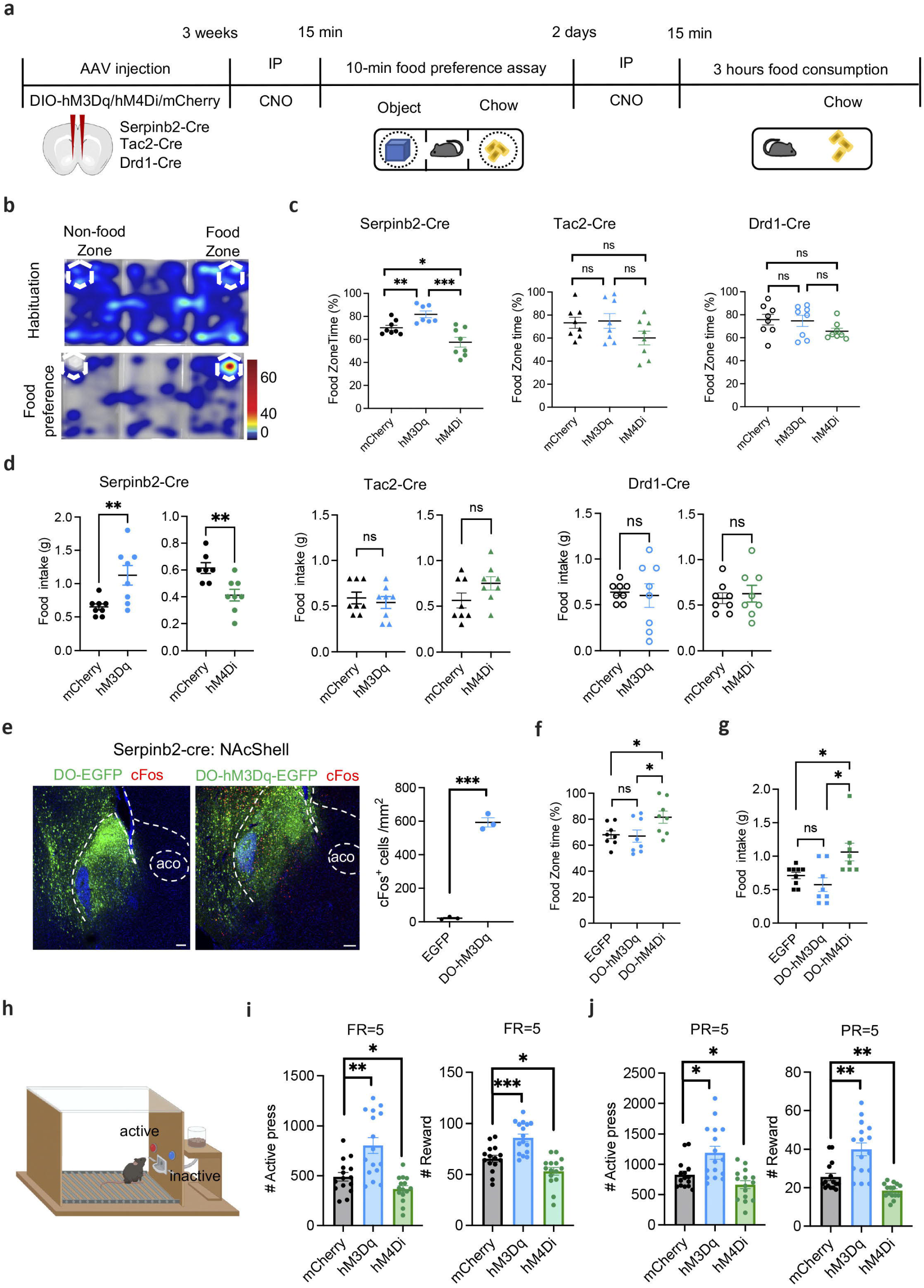
*Serpinb2*^+^ neurons bidirectionally regulate food seek and intake. **a,** Experimental scheme of the food approach and food consumption assays. **b,** Heat map encoding spatial location of a fasted mouse using the free access feeding paradigm. **c,** Percentage of time that mice spent in food zone. Chemogenetic activation (hM3Dq) or inhibition (hM4Di) of *Serpinb2*^+^ neurons (left), *Tac2*^+^ neurons (middle), and *Drd1*^+^ neurons (right). ***, p<0.001; **, p<0.01; *, p<0.05; ns, p>0.05, one-way ANOVA, with Tukey’s multiple comparisons. **d,** Total food consumption during the 3-hour test. Chemogenetic activation (hM3Dq) or inhibition (hM4Di) of *Serpinb2*^+^, *Tac2*^+^ and *Drd1*^+^ neurons. ***, p<0.001; **, p<0.01; *, p<0.05; ns, p>0.05, unpaired t-test. **e,** DO-EGFP/hM3Dq-EGFP virus expression in NAc Shell and number of *cFos^+^*neurons after CNO treatment. ***, p<0.001; **, p<0.01; *, p<0.05; ns, p>0.05, unpaired t-test. Scale bar: 100 μm. **f,** Percentage of time that mice spent in food zone. *, p<0.05; ns, p>0.05, one-way ANOVA test. **g,** Total food consumption during the 3-hour test. *, p<0.05; ns, p>0.05, one-way ANOVA test. **h,** Diagrammatic illustration of the food operant chamber paradigm. Mice were trained to press the lever to get food; pressing the active lever is followed by the delivery of food pellet, while pressing the inactive lever yields no outcome. The behavioral training includes habituation phase and fixed ratio (FR) training phase. Mice received CNO injection (2 mg/kg for the hM3Dq group and 5 mg/kg for the hM4Di group) 15 mins before they were placed into the operant chamber to start the FR and progressive ratio (PR) tests. **i,** Results of FR=5 test. The total number of active lever pressing (left) and total number of reward (right) after chemogenetic manipulations. ***, p<0.001; **, p<0.01; *, p<0.05; ns, p>0.05; one-way ANOVA test. **j,** Results of PR=5 test. The total number of active lever pressing (left) and total number of reward (right) after chemogenetic manipulations. ***, p<0.001; **, p<0.01; *, p<0.05; ns, p>0.05; one-way ANOVA test. Data are represented as mean ± SEM.

We further examined the involvement of the NAcSh *Drd1^+^* neurons in feeding by performing the experiments with the *Drd1*-Cre mice, and observed that the Ca^2+^ signal tended to increase before start eating the chow and gradually decrease after eating start with no significant changes were observed when sniffing an object (Fig. 2f1-f3). After ending chow eating, *Drd1^+^*neuronal activity increased concomitantly while Ca^2+^ signal showed random change upon interacting with an object (Fig. 2g1-g3). To quantify the *Drd1*^+^ neuronal activity at different events, we averaged the calcium signal of each animal and observed that the signals is significantly decreased after starting to eat chow (Fig. 2f4, left) but not when sniffing the object (Fig. 2f4, right). While the signals are significantly increased after ending chow eating (Fig. 2g4, left) but not when leaving the object (Fig. 2g4, right). These results indicate that *Drd1*^+^ neuronal activity increases when approaching the chow, decreases during eating and dramatically increases after eating ends. As expected, no obvious Ca^2+^ signals changes were captured for the eYFP injected *Serpinb2*-Cre or *Drd1*-Cre mice when mice eating or interacting with an object (Extended Data Fig. 3a2-3, c2-3).

Collectively, these results indicate that *Serpinb2*^+^ neurons respond to feeding particularly when eating start. On the other hand, the NAcSh *Drd1*^+^ neurons mainly respond when eating finishes. While *Tac2^+^* neurons do not respond to feeding.

### *Serpinb2*^+^ neurons bidirectionally regulate food intake

To determine whether *Serpinb2*^+^ neuronal activity plays a causal role in regulating feeding behavior, we examined whether the feeding behavior can be influenced by selectively manipulating the *Serpinb2*^+^ neuronal activity. To this end, we injected the Cre-dependent chemogenetic activation vector AAV-DIO-hM3Dq-mCherry or the inhibitory vector AAV-DIO-hM4Di-mCherry into the NAcSh region with AAV-DIO-mCherry vector as a control (Fig. 3a). We also performed parallel experiments using *Tac2*-Cre and *Drd1*-Cre mice (Extended Data Fig. 4a).

After confirming the injection accuracy (Extended Data Fig. 4b), and the efficiency of neuronal activation or inhibition with *cFos* expression (Extended Data Fig. 4c), we then tested whether activation or inhibition of the *Serpinb2*^+^ neurons affects feeding and reward-related behaviors (Fig. 3a). In the food approach assay, where mice have free access to 3 chambers, we analyzed the time mice spent in the food zone (Fig. 3b). As expected, mice from the control group spent significantly more time in the food zone than in the object zone. Importantly, activation (hM3Dq) or inhibition (hM4Di) of the *Serpinb2*^+^ neurons respectively increased or decreased the time the mice spent in the food zone compared to that of the control mice (Fig. 3c, left). In contrast, similar manipulations of the *Tac2^+^*or *Drd1^+^* neurons did not significantly alter the time the mice spent in the food zone compared to that of the control mice (Fig. 3c middle, right). Next, we measured the food consumption in home cage and found that activation of the *Serpinb2*^+^ neurons increased food consumption, while inhibition decreased food consumption (Fig. 3d, left). Similar manipulations performed on *Tac2*-Cre or *Drd1*-Cre mice did not affect food consumption (Fig. 3d middle, right).

Since our Ca^2+^ imaging results suggested that *Serpinb2*^+^ neuronal activity is positively associated with eating start while NAcSh *Drd1*^+^ neuronal activity decreases upon eating, we hypothesized that *Serpinb2*^-^ and *Serpinb2*^+^ neurons play opposite roles in regulating feeding. To test this hypothesis, we used a Cre-off virus (Double-floxed Orientation, DO) to label all neurons that do not express Cre recombinase, and examined the *Serpinb2*^-^ neurons in regulating feeding. To this end, we injected *Serpinb2*-Cre mice with DO-hM3Dq-EGFP/hM4Di-EGFP for activation/inactivation with DO-EGFP serves as a control. Virus expression pattern and activation effect were confirmed by immunostaining (Fig. 3e). We then performed the food approach, and food consumption tests on the Cre-OFF virus injected mice. Inactivation of *Serpinb2*^-^ neurons induced a significant increase in the time spent in the food zone with concomitant increase in food consumption (Fig. 3f, g), which is opposite to the manipulation of the *Serpinb2*^+^ neurons (Fig. 3c, d). These results indicate that *Serpinb2*^+^ neurons have a specific role in feeding that could be counteracted by other *Serpinb2*^-^ neurons in NAcSh.

Since the NAcSh functions as a hub modulating motivations, we next carried out the food operant chamber test to further determine whether the neuronal activity of the *Serpinb2*^+^ neurons has a causal role in regulating food motivation (Fig. 3h). To maintain a similar food motivation status, all mice were food restricted to reduce body weight to around 90% of their original value. After trained to operantly respond to chow pellets on fixed ratio (FR) 1, 3, 5 schedule, the animals were then tested for lever pressing upon CNO-induced chemogenetic manipulation. For the FR 5 test, activation of the *Serpinb2*^+^ neurons significantly increased the active lever pressing and pellet reward (Fig. 3i), while inhibition elicited the opposite effect (Fig. 3i). A similar result was also obtained when a progressive ratio (PR) 5 was used for the test (Fig. 3j). Taken together, these results demonstrate that the *Serpinb2*^+^ neurons are involved in bidirectionally control of the goal-direct food seeking behavior.

In addition to food consumption, the NAcSh has been shown to participate in other motivated behaviors, including conditioned reinforcement^38, 39^, hedonic reactivity^40^, anxiety^41^ and social play^42^. Thus, we asked whether *Serpinb2*^+^ neurons also regulate these behaviors under the same chemogenetic manipulation. We found that manipulation of *Serpinb2*^+^ neuronal activity does not affect locomotion in open field test (Extended Data Fig. 5a, b), conditioned place preference (CPP) (Extended Data Fig. 5c), cocaine CPP test (Extended Data Fig. 5d), anhedonia in sucrose preference test (Extended Data Fig. 5e), social interaction (Extended Data Fig. 5f, g) or anxiety in the elevated plus maze test (Extended Data Fig. 5h, i). These results support that *Serpinb2*^+^ neurons are specifically involved in feeding, but not in locomotion, anxiety, social, anhedonia or drug seeking behaviors.

### *Serpinb2*^+^ neurons mediate food consumption via LH projection

Thus far, we have demonstrated a critical role of *Serpinb2*^+^ neurons in regulating feeding behaviors. Next, we attempted to reveal the circuit mechanism underlying the functions of *Serpinb2*^+^ neurons in regulating food consumption. Previous studies have indicated that NAcSh D1-MSNs project to multiple brain regions, including the ventral tegmental area (VTA)^43^, LH^20^and ventral pallidum (VP)^44^. To determine the projection sites of *Serpinb2*^+^ neurons, we injected Cre-dependent AAVs expressing membrane-bound GFP (mGFP, for labeling axons) and synaptophysin-mRuby (SYP-mRuby, for labeling putative presynaptic sites) into the NAcSh of the *Serpinb2*-Cre mice (Fig. 4a). Three weeks after the virus injection, we analyzed the brain sections and observed colocalization of green neuronal terminals and red pre-synaptic puncta only in the lateral hypothalamus (LH) (Fig. 4b, c), but not in VP, BLA or VTA (Extended Data Fig. 6), indicating that the *Serpinb2*^+^ neurons only project to LH. To further validate this projection, we injected the retrograde tracer Cholera toxin subunit B conjugated with Alexa Fluor 647 (CTB-647)^45^ into the LH region (Fig. 4d) and DIO-mCherry into the NAcSh of *Serpinb2*-Cre mice. Immunostaining showed colocalization of CTB with mCherry in the NAcSh (Fig. 4e), supporting the notion that the NAcSh *Serpinb2*^+^ neurons are project to the LH.

**Fig. 4.**
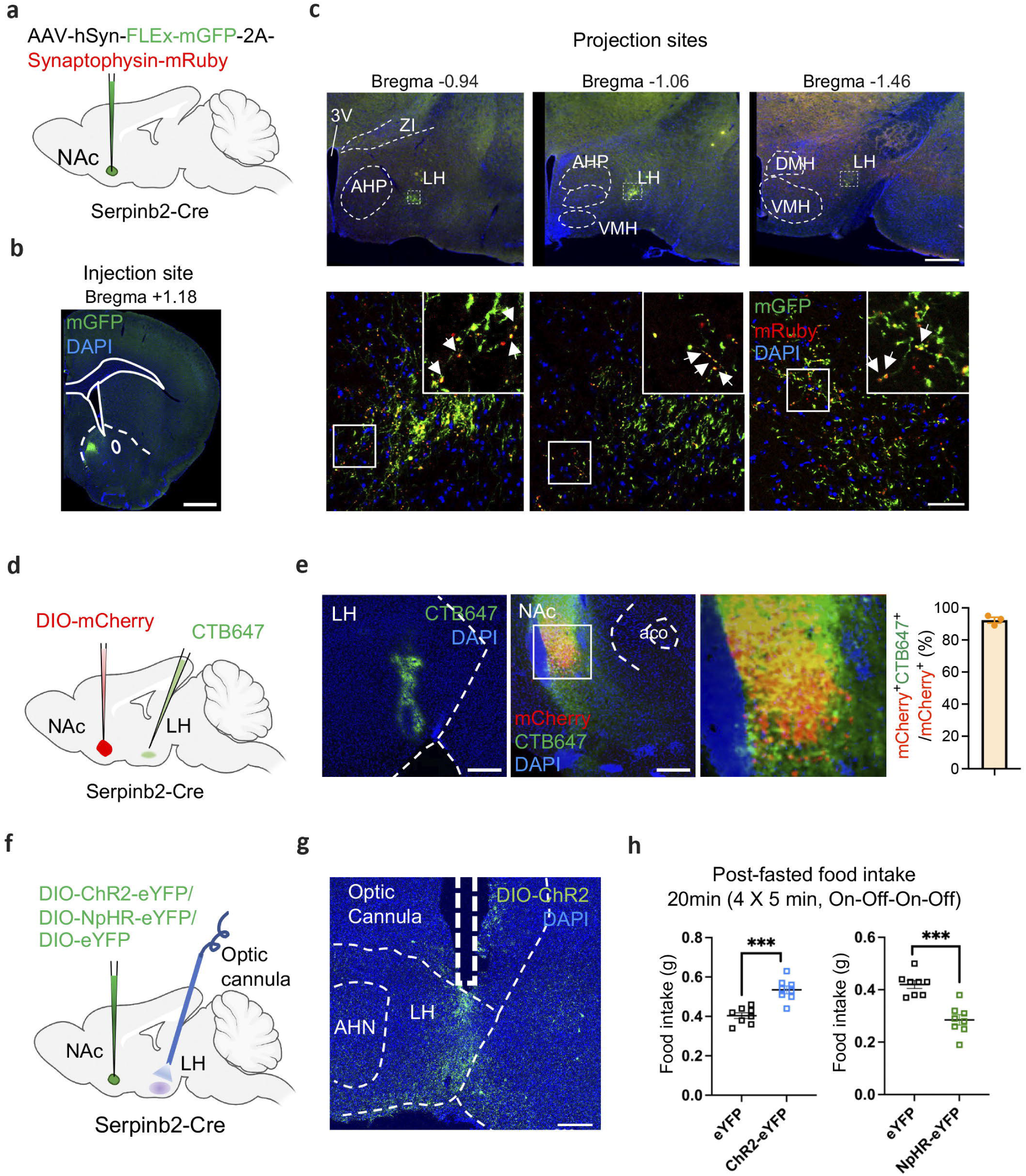
*Serpinb2*^+^ neurons mediate food intake via LH projection. **a, b,** Diagram of injection of the NAc *Serpinb2*^+^ neurons with AAV-hSyn-FLEx-mGFP-2A-Synaptophysin-mRuby (**a**), the expression FLEx-mGFP in the NAcSh (**b**). Scale bar, 100 μm. **c,** Top, *Serpinb2*^+^ neuron projection to LH. Scale bar: 500 μm. Bottom, enlarged image of the square. Scale bar: 50 μm. **d,** Diagram of retrograde tracing approach. **e,** Left, validation of injection site of CTB 647. Middle, validation of injection site of DIO-mCherry in NAcSh. Right, colocalization of retrograde tracing of LH neurons with CTB 647, with the *Serpinb2*^+^ neurons labeled by DIO-mCherry. Scale bar: 50 μm. **f, g,** Diagram illustrate the indicated AAV injection into NAc and optic cannulas implantation in LH area (**f**), and histology validating virus expression and cannula implantation site (**g**). Scale bar: 200 μm. **h,** Optogenetic activation (left) or inhibition (right) of NAc *^Serpinb2^*^+^→LH neurons respectively increased or decreased food intake. Data in (**h)** is presented as mean ± SEM. ***, p<0.001; **, p<0.01; *, p<0.05; ns, p>0.05, unpaired t-test. 3V, third ventricle; AHN, anterior hypothalamic nucleus; AHP, anterior hypothalamic area, posterior part; DMH, ventromedial hypothalamic nucleus; HY, hypothalamus; VMH, ventromedial hypothalamic nucleus; ZI, Zona incerta.

To demonstrate that the LH projection of *Serpinb2*^+^ neurons is functionally relevant to feeding, we asked whether optogenetic manipulation of the *Serpinb2*^+^ neuron terminals in LH can regulate the feeding behavior. To this end, AAV-DIO-ChR2-eYFP or AAV-DIO-NpHR-eYFP vectors were injected to the NAcSh of the *Serpinb2* -Cre mice with optic cannula implanted into their LH region, while AAV-DIO-eYFP vector was used as a control (Fig. 4f, g). After fasting the mice for overnight, we activated the *Serpinb2*^+^ neuron terminals in the LH with blue light on-off stimulation (20 Hz, 2-ms pulses). We found that stimulation of *Serpinb2*^+^ neuron terminals in ChR2 expressing mice significantly increased the food intake compared to the control mice that express eYFP (Fig. 4h, left panel). Conversely, inhibition of the *Serpinb2*^+^ neuron terminal in the LH with yellow light decreased total food intake compared to the control (Fig. 4h, right panel). Collectively, viral tracing and circuit manipulation demonstrate that the NAcSh *Serpinb2^+^* neurons to LH projection is functionally important for food intaking behaviors.

### *Serpinb2*^+^ neurons form a circuit with LH LepR^+^ GABA^+^ neurons and counterbalance the effects of leptin

Having demonstrated the functional importance of the NAcSh to LH projection, we next attempted to determine the neuron types in LH that receive signals from the *Serpinb2*^+^ neurons. The LH is a highly heterogeneous brain region controlling food intake, energy expenditure, and many other physiological functions^46^. Since neuropeptides orexin/hypocretin and melanin-concentrating hormone (MCH) are associated with feeding^47, 48^, and are mainly expressed in the LH, we first asked whether they are the down-stream targets of the NAcSh *Serpinb2*^+^ neurons. To this end, we injected the AAV-hSyn-FLEx-mGFP-2A-Synaptophysin-mRuby viruses to the NAcSh of the *Serpinb2* -Cre mice and performed immunostaining of candidate neuropeptides or transmitters on slices covering the LH (Fig. 5a). We found that no MCH- or orexin-expressing neurons in the LH (Fig. 5b, c, indicated by arrows) overlapped with *Serpinb2*^+^ (green) terminals. Given that GABAergic neurons are the most abundant subtype in the LH and are involved in feeding and leptin-regulated energy homeostasis^11, 49, 50^, we next checked whether leptin receptor (LepR) positive GABAergic neurons^19^ receive projections from NAcSh *Serpinb2^+^* neurons. After immunostained with anti-GABA or anti-LepR antibodies, we found that the presynaptic puncta of *Serpinb2^+^* neuron terminals were located around the LepR^+^ GABA^+^ somas (Fig. 5d, e, arrow heads). To confirm the connection of NAcSh *Serpinb2^+^* neurons and LH *LepR*^+^ neurons, we performed monosynaptic rabies virus tracing by injecting Cre-dependent AAVs expressing mCherry-TVA and rabies G protein into the LH of LepR-cre mice. Two weeks later, Enva-ΔG rabies virus expressing GFP was injected into the same location, which revealed NAc as one of the inputs for LH^LepR+^ neurons (Fig. 5f, g). smFISH further confirmed that the GFP labeled neurons are Serpinb2-expressing neurons (Fig. 5g), which supports that NAcSh *Serpinb2^+^* neurons directly target LH^LepR+^ neurons. To demonstrate that the NAcSh^Serpinb2+^-LH^LepR+^ circuit is functional, we selectively activated or inhibited *Serpinb2^+^* neurons and then analyzed the LH^LepR+^ neuronal response. Since MSNs in NAc are GABAergic neurons, inhibition of *Serpinb2^+^*neurons would lead to decreased inhibition effect on the downstream neurons. As expected, we observed increased *cFos* signals which merged with LepR signal in LH (Fig. 5i, right bottom panel). Importantly, inhibition of *Serpinb2^+^*neurons significantly increased the cFos^+^ neurons in LH compared to that in the control (mCherry) or activation (hM3Dq) groups (Fig. 5j). Thus, NAcSh^Serpinb2+^ neurons are functionally connected with LH^LepR+^ neurons.

**Fig. 5.**
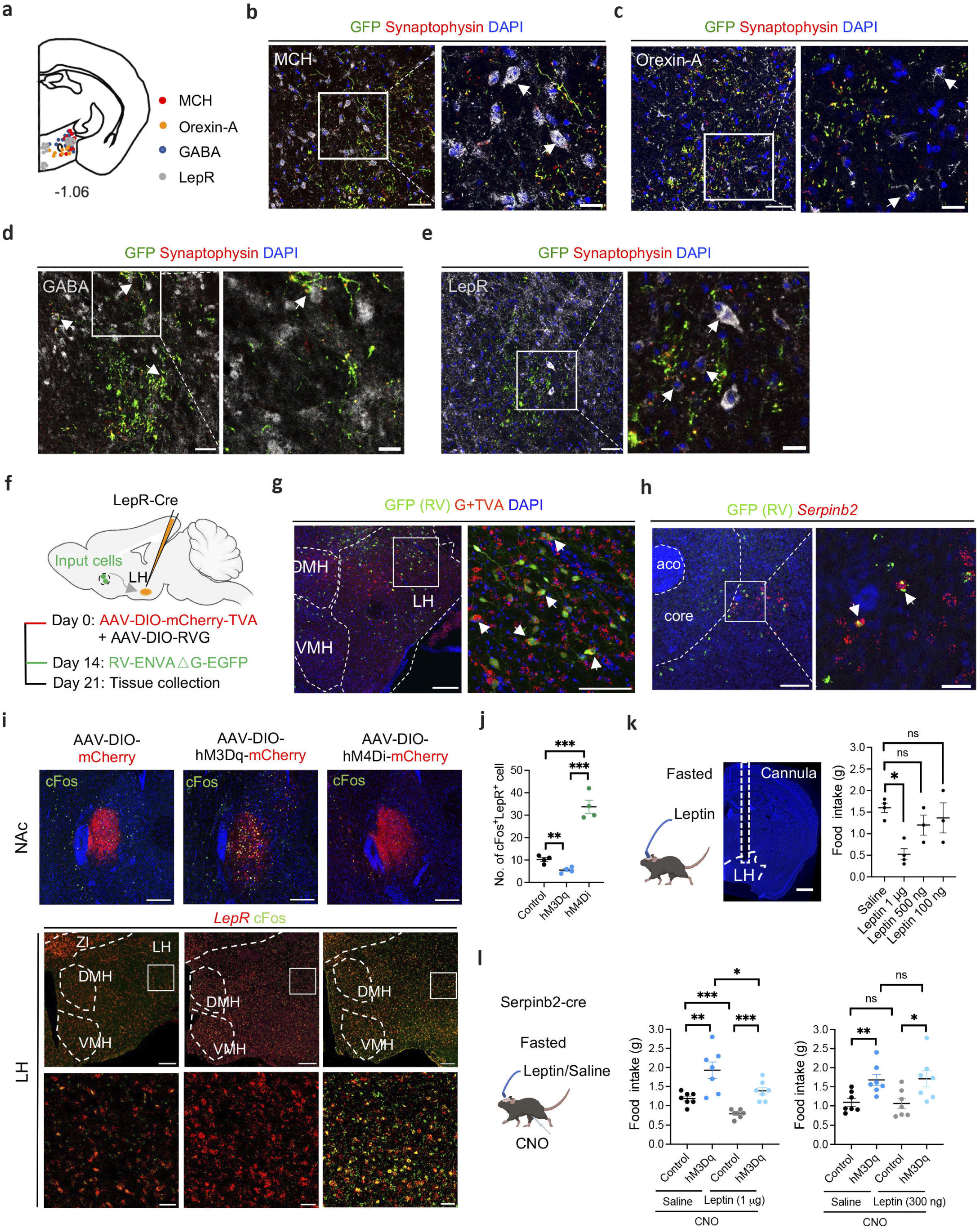
*Serpinb2*^+^ neurons project to LH LepR^+^ GABA^+^ neurons and their activation can counteract leptin effect. **a,** Diagram indicating the MCH^+^, Orexin-A^+^, GABA^+^ and LepR^+^ neurons in LH. **b,** Example images of not-connected MCH^+^ neurons (grey) and *Serpinb2*^+^ neuron terminals (green fibers with red puncta) in LH. Scale bar, left, 50 μm; right, 20 μm. Arrows indicate somas. **c,** Example images of not-connected Orexin-A^+^ neurons (grey) and *Serpinb2*^+^ neuron terminals (green fibers with red puncta) in LH. Scale bar, 50 μm; right, 20 μm. Arrows indicate somas. **d,** Images showing the co-localization of *Serpinb2*^+^ neuron terminals (green fibers with red puncta) with GABA^+^ (grey) as indicated by arrows. Scale bar, left, 50 μm, right, 20 μm. **e,** Images showing the co-localization of *Serpinb2*^+^ neuron terminals (green fibers with red puncta) with LepR^+^ neurons (grey) as indicated by arrows. Scale bar, left, 50 μm, right, 20 μm. **f,** The timeline of monosynaptic retrograde rabies tracing of LH^LepR^ neurons. **g,** Representative images showing the location of starter LH^LepR^ neurons in a LepR-cre mouse. TVA-mCherry (red), Rabies-GFP (green) and DAPI (blue), scale bars, 100 µm (left), 50 µm (right). **h,** Representative histological images with cells retrogradely labeled from LH^LepR^ neurons (green) and *Serpinb2*^+^ neurons (red). Scale bar, 200 µm (left) and 50 µm (right). **i,** cFos detection in LH of DREAADs treated *Serpinb2*-cre mice. Virus expression in NAcSh (top). mCherry (red), cFos (green) and DAPI (blue). CNO-induced cFos immunofluorescence signals were observed in LH, bottoms are enlarged images of the square. Scale bar: 100 µm (top), 200 µm (middle) and 50 µm (bottom). **j,** The number of cFos^+^ cells were compared among the three groups (n=4 sections from 4 mice). ***, p<0.001; **, p<0.01; *, p<0.05; ns, p>0.05; one-way ANOVA test. **k,** Diagram showing cannula implantation in LH for leptin delivery (left, middle). CNO delivery was achieved via i.p. injection. Results of total food consumption in 3 hours by fasted mice with different doses of leptin administration (right). Scale bar: 500 μm. ***, p<0.001; **, p<0.01; *, p<0.05; ns, p>0.05; one-way ANOVA test. **l,** Same as panel K except food consumption is quantified under different conditions with or without *Serpinb2^+^*neuron activation in the presence or absence of 1 μg (middle) or 300 ng (right) of leptin delivery. ***, p<0.001; **, p<0.01; *, p<0.05; ns, p>0.05; one-way ANOVA test. Data is presented as mean ± SEM.

As an adipose-derived hormone, leptin plays a central role in regulating energy homeostasis^51–53^. Leptin performs most of its functions, including suppression of food intaking, by activating the LepR on central nerve system (CNS) neurons^54, 55^. Since the NAcSh *Serpinb2*^+^ neurons project to LepR^+^ GABAergic neurons in LH (Fig. 5e-j), we anticipate that both leptin and the NAcSh *Serpinb2*^+^ neurons have shared neuron targets and consequently they should have functional interaction. To test for their potential functional interaction in food intake, we implanted a catheter in the LH for leptin delivery (catheter administration) in the *Serpinb2-*Cre mice that were also injected with hM3Dq-mCherry-expressing AAV into the NAcSh so that the NAcSh *Serpinb2*^+^ neurons can be activated by CNO via i.p. injection. After establishing that 1 μg of bilateral intra-LH leptin cannula delivery^56^ can significantly decrease food intake in 3 hours relative to the saline control (Fig. 5k), we used 1 μg of leptin for all subsequent tests. Consistent with what we have shown (Fig. 3d), CNO-induced *Serpinb2*^+^ neuron activation increased food intake (Fig. 5l, middle, control vs hM3Dq in Saline). Importantly, despite leptin delivery reduced food intake, 1 μg of leptin’s effect can be at least partly overcome by CNO-induced *Serpinb2*^+^ neuron activation (Fig. 5l, middle, hM3Dq with or without Leptin). While lower dose of leptin (300 ng) did not exhibit a similar effect (Fig. 5i, right). These results indicate that leptin-induced inhibitory effect on food intake can be at least partly overcome by *Serpinb2*^+^ neuron activation.

### Ablation of *Serpinb2*^+^ neurons promote weight loss and increase metabolic level

To assess whether loss function of the *Serpinb2*^+^ neurons can exert a long-term effect on energy homeostasis, we selectively ablated NAcSh *Serpinb2^+^*neurons in *Serpinb2-*Cre mice by injecting a Cre-dependent AAV vector expressing caspase-3 which eliminates the neurons by inducing cell death (Fig. 6a). Ablation of the NAcSh *Serpinb2*^+^ neurons resulted in decreased food intake (Fig. 6b) as well as reduced body weight gain (∼10% or 2g over 7 weeks) despite food and water being freely available in the home cage (Fig. 6c). A reduction on body weight is normally caused by an imbalance between energy intake and expenditure. To this end, we analyzed the effect of *Serpinb2^+^*neuron ablation on metabolism by performing metabolism recording, which measures the volumes of oxygen consumption (VO_2_), carbon dioxide production (VCO_2_), respiratory exchange ratio (RER), energy expenditure (XTOT), ambulatory movement (XAMB) and food intake (Fig. 6d-g, Extended Data Fig. 7a-c). We found that *Serpinb2^+^*neuron ablation (the taCasp3 group) significantly increased O_2_ consumption (Fig. 6d), VCO_2_ production (Fig. 6e), energy expenditure (Fig. 6f) as well as decreased cumulative food intake (Fig. 6g). However, no significant change in RER or movement was observed (Extended Data Fig. 7a-c). Given that a previous study indicated that the NAcSh to LH could modulate energy expernditutre^57^, the metabolic changes observed could be a result of lack of innervation from *Serpinb2*^+^ neurons. Collectively, these results demonstrate that ablation of *Serpinb2*^+^ neurons not only reduces food intake, but also increases energy expenditure, both can contribute to long-term bodyweight loss.

**Fig. 6.**
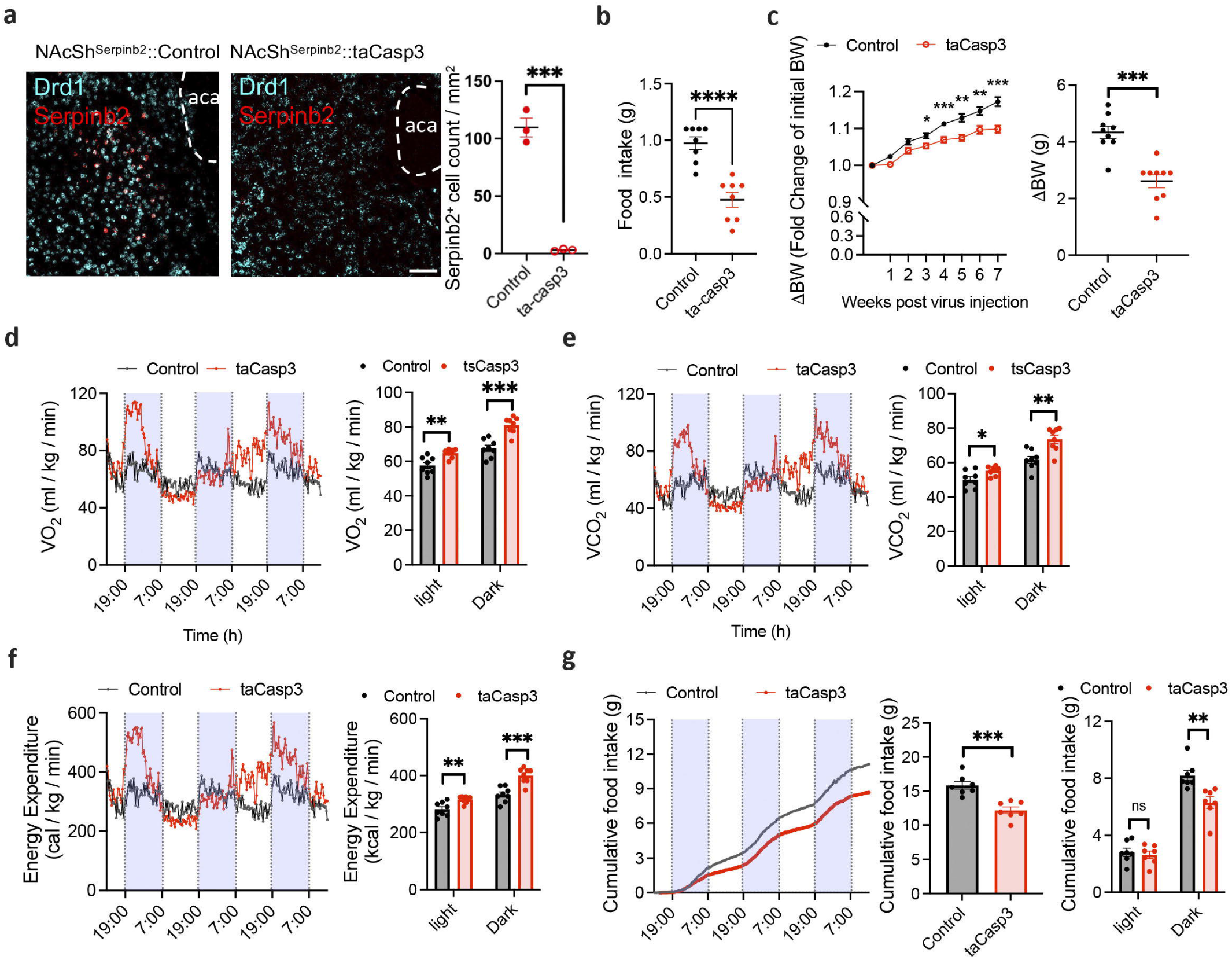
Ablation of *Serpinb2*^+^ neurons decrease bodyweight gain via reducing food consumption and increasing energy expenditure. **a,** FISH and quantification verify *Serpinb2^+^* neuron ablation after AAV-DIO-taCasp3 injection. Scale bar: 100 μm. ***P ≤ 0.001, ns, p>0.05, unpaired t-test. **b,** *Serpinb2^+^* neuron ablation decreased 3 hours total food consumption by mice. ***P ≤ 0.001, ns, p>0.05, unpaired t-test. **c,** *Serpinb2^+^* neuron ablation has a long-time effect on bodyweight loss. ***P ≤ 0.001, **P ≤ 0.01, *P ≤ 0.05; ns, P > 0.05, two-way ANOVA, Sidak’s test (left); unpaired t-test (right). **d,** Oxygen consumed (VO2) by GFP-expressing control (n = 8) and taCasp3 (n = 8) mice fed chow. ***, p<0.001; **, p<0.01; *, p<0.05; ns, p>0.05; unpaired t-test. **e,** Carbon dioxide consumed (VO2) by GFP-expressing control (n = 8) and taCasp3 (n = 8) mice fed chow. **P ≤ 0.01, *P ≤ 0.05; ns, unpaired t-test. **f,** Energy expenditure of mice by GFP-expressing control (n = 8) and taCasp3 (n = 8) mice fed chow. ***, p<0.001; **, p<0.01; *, p<0.05; ns, p>0.05; unpaired t-test. **g,** Cumulative food consumption from baseline to day 3. GFP-expressing control (n = 8) and taCasp3 (n = 8) mice fed chow. ***P ≤ 0.001, **P ≤ 0.01, *P ≤ 0.05; ns, P > 0.05, unpaired t-test.

## Discussion

Feeding is an essential goal-directed behavior that is heavily influenced by homeostatic state and motivation. The accumbal-to-lateral hypothalamic pathway has been implicated in regulating feeding behavior, but the specific neuron subtypes and precise neuronal circuit in LH are not clear. In this study, we filled in this knowledge gap by identifying a NAcSh D1 neuronal subtype and the associated circuit that integrates neuronal and humoral signal to regulate food consumption. Specifically, we identified a *Serpinb2-*expressing D1-MSN subtype located in the NAcSh that regulates the feeding behavior through the projection to LH^LepR^ neurons, providing innervations to LH^LepR^ neurons from outside the hypothalamus. We demonstrate that the *Serpinb2*^+^ neurons bidirectionally modulate food motivation and consumption specifically without affecting other motivated behaviors, such as cocaine CPP, sucrose preference, or social interaction. Importantly, *Serpinb2*^+^ neurons target the LepR-expressing GABAergic neurons in LH and their activation can partly overcome the suppressive effect of leptin on food intaking, while their ablation leads to increased energy expenditure, and decreased food consumption, cumulating bodyweight loss.

### The *Serpinb2^+^* neurons are functionally distinct from the pan D1-MSNs in NAcSh

Previous studies have observed reduced activity of D1-MSNs during food consumption, and consistently, suppressing D1-MSNs activity prolonged food intake^20^. Using *Serpinb2*-Cre, *Tac2*-Cre and *Drd1*-Cre mice, we compared the *Serpinb2^+^*, *Tac2*^+^ and the *Drd1*^+^ MSNs in regulating feeding behaviors, and found manipulations of these neuron subtypes led to different outcomes (Figs. 2 and 3). First, *Serpinb2^+^* neurons are activated by food consumption but suppressed after ending eating, while *Drd1^+^* neurons show a minor activation during food approaching but a greater activation after food consumption. Second, *Serpinb2^+^* neurons bidirectionally regulate food seeking and intake, particularly during refeeding, while manipulating *Drd1^+^* neurons do not significantly alter feeding behavior. On the other hand, a previous study showed that inhibition of *Drd1^+^*neurons promoted liquid fat food intake^20^. The difference might be due to the different feeding assays used in the two studies. We used free-access food intake in this study, while the previous study used a head-fixed mice licking liquid fat food as the assay^20^. Third, ablation of the *Serpinb2^+^*neurons significantly reduced food intake (Fig. 6), which is consistent with our finding that *Serpinb2^+^* neuron activation positively regulates food intake (Fig. 3). However, a previous study indicated that lesions or inactivation of the NAc neurons do not significantly alter food consumption^58^. We do not consider these results to be in conflict as NAc is composed of many D1- and D2-MSN neuron subtypes, of which many may not involve in regulating food intake, while others can positively or negatively regulate food intake. Consequently, manipulating *Serpinb2^+^*neurons and the entire NAc neurons can have different outcomes. Indeed, using genetically engineered Cre-off virus, we demonstrated that *Serpinb2^+^*neurons and *Serpinb2^-^* neurons have opposite roles in regulating feeding (Fig. 3c,d compared to f, g). Our results indicate that finer granularity and cell type-specific approaches are needed to dissect the function of different neuron subtypes in NAc. For example, although the *D2^+^*neuronal activity as a whole is not altered during food consumption^20, 28^, the D2 receptors are indeed downregulated in obese rodent and humans^59, 60^. Whether certain D2-MSN subtypes are involved in regulating food intake remains to be determined.

### *Serpinb2^+^* and *Tac2^+^* MSNs respectively modulate food and drug reward

The NAcSh has long been implicated in regulating reward-related behaviors that are associated with food, social, and drug. Previous studies were mainly based on dichotomous MSNs subtypes that express dopamine receptor 1 or dopamine receptor 2 (D1- or D2-MSNs)^61^. Increasing evidence suggest that the NAcSh is highly heterogeneous in terms of the molecular features and anatomical connections of the neurons located in this region. This raises an interesting question that whether these reward-related behaviors are regulated by distinct or overlap neuron subtypes and/or projections. By combining MERFISH analysis, *cFos* mapping, neuronal activity manipulation of different subtypes with behavioral tests, we found that the *Serpinb2^+^* D1-MSNs of NAcSh specifically regulate food reward, but not drug reward or other emotional and cognitive functions (Fig. 1, 3, Extended Data Fig. 5). On the other hand, we have previously showed that the NAcSh *Tac2^+^* D1-MSNs specifically regulate cocaine reward^62^. These studies indicate that different reward behaviors are at least partly regulated by distinct MSNs subtypes. An important task in future studies will be to identify the relevant neuron subtypes regulating the various reward-related behaviors, and eventually to link the cellular heterogeneity to functional diversity of each brain regions. This way, the cellular and circuit mechanisms underlying the various behaviors can be elucidated.

### The NAcSh*^Serpinb2^*^+^ -LH*^LepR^*^+^ circuit controls feeding in hungry state

Previous studies have shown that either LH GABA or Vglut2 neuronal subpopulation can receive NAc innervation^20, 63^. However, the LH GABA and Vglut2 neurons are extremely heterogeneous, and can be further divided into 15 distinct populations, respectively^64^. Thus, the specific cell types that receive NAc innervation were unknown. Previous studies also showed that NAcSh D1-MSN to LH inhibitory transmission stops eating, and endocannabinoids mediated suppression of this projection promotes excessive eating of highly palatable chow^65^, but the D1 subtype involved in this projection was not known. Using viral tracing, we discovered that the NAcSh *Serpinb2*^+^ D1-MSNs project to *LepR*^+^ neurons in LH underlying the *Serpinb2*^+^ neuron function in food intake (Fig. 5e-j). Distributed in numerous regions involved in the regulation of energy balance, the *LepR*^+^ neurons lie in the mediobasal hypothalamic (MBH) “satiety centers” and in LH that is regarded as the “feeding center”^53, 66^. Leptin treatment induced *cFos* expression and 100 nM of leptin depolarized 34% of LepR-expressing neurons in LH^19^. Unilateral intra-LH leptin decreased food intake and bodyweight^19^. In our study, we found activation of the *Serpinb2*^+^ neurons increased the inhibition of the *LepR*^+^ neurons excitability, resulting in increased food consumption; while inhibition of *Serpinb2*^+^ neurons decreased the inhibition of *LepR*^+^ neurons excitability, leading to decreased food consumption even after fasting (Fig. 3d, 4h). Our results are consistent with previous reports demonstrating that LH LepR neuron activation decreases chow intake^67^. Importantly, manipulating *Serpinb2*^+^ neuronal activity could at least partly override leptin’s effect in LH to modulate food consumption (Fig. 5l). Our study thus reveals a parallel and compensatory circuit, from NAcSh to LH^LepR^, beyond the hypothalamus circuit that directly modulates food intake, to maintain energy homeostasis.

### Regulation of energy homeostasis beyond hypothalamus

Feeding is an essential goal-directed behavior that is influenced by cellular homeostasis state and appetitive motivation. Given the extensive nature of the NAcSh-to-LH projection, its function likely extends beyond regulating food consumption. In our study, we conducted operant food intake assay and found that *Serpinb2^+^* neuronal activity bi-directionally regulates active lever presses and the earned reward, which reflect appetitive food motivation (Fig. 3i, j). A role of *Serpinb2^+^* neurons in regulating energy homeostasis and appetitive motivation is consistent with previous finding that NAc is involved in integrating descending signals pertaining to homeostatic needs and goal-related behaviors^5, 68^. On the other hand, ablation of the *Serpinb2^+^* neurons leads to bodyweight loss, which is a combined effect of reduced food intake and increased energy expenditure. These observations raise the possibility that the NAcSh^Serpinb2+^ to LH^LepR+^ projection may provide a regulatory mechanism that enable rapid switching between different behavioral states in response to changing external conditions. Clinically, activation of the NAcSh *Serpinb2^+^* neurons may provide a potential strategy for anorexia or obese treatment.

In conclusion, by focusing on the neuron subtypes located in the NAcSh, we identified a molecularly defined neuron subtype that can regulate food intake through a neuron-hormone axis. Our detailed characterization of the *Serpinb2^+^* neuron subtype not only clarified previous discrepancy regarding the role of NAcSh to LH circuit in regulating feeding behavior, but also revealed *LepR*^+^ neurons in LH as the downstream neurons receiving the inputs, thus linking neuronal and hormonal regulation. In addition, our findings have clinical implication as activating or ablating a small *Serpinb2^+^*neurons in NAcSh could regulate food intake and metabolism. Thus, the small population of NAcSh *Serpinb2^+^* neurons could be an ideal entry point for understanding the complex brain-metabolism regulatory network underlying eating and bodyweight control.

## Methods

### Animals

All experiments were conducted in accordance with the National Institute of Health Guide for Care and Use of Laboratory Animals and approved by the Institutional Animal Care and Use Committee (IACUC) of Boston Children’s Hospital and Harvard Medical School. The *Serpinb2*-Cre mice were generated by Cyagen US Inc. The *Tac2*-Cre knock-in mouse line was a gift from Q. Ma at Dana-Farber Cancer Institute and Harvard Medical School. 129-Tg (Drd1-cre)120Mxu/Mmjax mice (Jax: 037156), B6.Cg-Gt(ROSA)26Sor^tm9(CAG-tdTomato)Hze^/J (Jax:007909) and C57BL/6NJ (Jax:000664) mice were purchased from Jackson lab. For behavioral assays, 12-16 weeks old male mice were used. The mice were housed in groups (3-5 mice/cage) in a 12-hr light/dark cycle (light time, 7:00 to 19:00), with food and water *ad libitum* unless otherwise specified.

### Fluorescence *in situ* hybridization (FISH) and immunofluorescence (IF) staining

Mice were transcardially perfused with PBS followed by 4% paraformaldehyde. Brains were then placed in a 30% sucrose solution for 2 days. The brains were frozen in Optimal Cutting Temperature (OCT) embedding media and 16 μm (for FISH) or 35 μm (for IF) coronal sections were cut with vibratome (Leica, no. CM3050 S). For FISH experiments, the slices were mounted on SuperFrost Plus slides, and air dried. The multi-color FISH experiments were performed following the instructions of RNAscope Fluorescent Multiplex Assay (ACD Bioscience). The probes used in this study listed as bellow: Tac2 (ACD, Cat No. 446391-C2), Serpinb2 (Cat No. 538091), Drd1(Cat No. 461901-C3), Drd2 (Cat No. 406501-C3), Spon1 (Cat No. 492671), Stard5 (Cat No. 880931-C3) and Lepr (Cat No. 402731-C3). For IF, cryostat sections were collected and incubated overnight with blocking solution (1×PBS containing 5% goat serum, 5% BSA, and 0.1% Triton X-100), and then incubated with the following primary antibodies, diluted with blocking solution, for 1 day at 4 °C: rabbit anti-cFos (1:2000, Synaptic systems, #226003), chicken anti-GFP (1:2000, Aves Labs, no. GFP-1010), chicken anti-mCherry (1:2000, Novus Biologicals, no. NBP2-25158), mouse anti-Orexin-A(KK09) (1:500, Santa Cruz Biotechnology, Cat#sc-80263), rabbit anti-GABA (1:1000, Sigma, Cat#A2052), rabbit anti-MCH (1:20000, Phoenix Pharmaceuticals, Cat#H-070-47), rabbit anti-Leptin receptor (1:1000, Abcam, Cat#104403). Samples were then washed three times with washing buffer (1×PBS containing 0.1% Tween-20) and incubated with the Alexa Fluor conjugated secondary antibodies (1:500, Thermo Fisher Scientific, Cat#A11039, 21206,10042) for 2 h at room temperature. The sections were mounted and imaged using a Zeiss LSM800 confocal microscope or an Olympus VS120 Slide Scanning System.

### AAV vectors

The following AAV vectors (with a titer of >10^12^) were purchased from UNC Vector Core: AAV5-EF1a-DIO-hChR2(H134R)-EYFP, AAV5-EF1a-DIO-EYFP, AAV-DJ-EF1a-DIO-GCaMP7s. The following AAV vectors were purchased from Addgene: AAV5-hSyn-DIO-hM3D(Gq)-mCherry (#44361), AAV5-hSyn-DIO-hM4D(Gi)-mCherry (#44362), AAV5-hSyn-DIO-mCherry (#50459), pAAV-flex-taCasp3-TEVp (#45580), pAAV-Ef1α-DIO eNpHR 3.0-EYFP (#26966), pAAV-hSyn-FLEx-mGFP-2A-Synaptophysin-mRuby (#71760). The following AAV vectors were purchased from BrainVTA: rAAV-hSyn-DO-hM3D(Gq)-EGFP-WPREs (PT-2155), rAAV-hSyn-DO-hM4D(Gi)-EGFP-WPREs (PT-6701), rAAV-hSyn-DO-EGFP-WPREs (NA), rAAV-Ef1a-DIO-mCherry-F2A-TVA-WPRE-hGH-polyA (PT0023), rAAV-EF1a-DIO-RVG (PT0207) and RV-ENVA-ΔG-EGFP (R01001).

### Stereotaxic brain surgeries

The AAV vectors were injected through a pulled-glass pipette and the nanoliter injector (Nanoject III, Drummond Scientific −3-000-207). The injection was performed using a small-animal stereotaxic instrument (David Kopf Instruments, model 940) under general anesthesia by isoflurane (0.8 liter/min, isoflurane concentration 1.5%) in oxygen. A feedback heater was used to keep mice warm during surgeries. Mice were allowed to recover in a warm blanket before they were transferred to housing cages for 2–4 weeks before behavioral evaluation was performed. For chemogenetics experiments, 0.1∼0.15 μl of AAV vector was bilaterally delivered into target regions. For optogenetics experiments, following viral injection, the fiber optic cannula (200 μm in diameter, Inper Inc.) were implanted 0.1 mm above viral injection site and were secured with dental cement (Parkell, #S380). For the drug delivery cannula implantation, the cannula (Guide cannula: C.C 2.0mm, C=4.5mm; Injector: G1=0.5mm; Dummy cannula: G2=0, RWD Life Science) were directly implanted 0.5 mm above the LH and were secured with dental cement (Parkell, #S380). The coordinates of viral injection sites are based on previous literature as follow: NAc (Anterior-Posterior [AP] +1.2, Medial-Lateral [ML] ± 0.6, Dorsal-Ventral [DV] −4.5 mm) and LH ([AP] −1.3, [ML] ± 1.2, [DV] −5.0 mm).

### Neuronal Tracing

For CTB tracing, mice were injected with 0.1-0.2 μl CTB-647 (AF-CTB, all from Life Technologies) unilaterally into the LH ([AP] −1.2, [ML] +1.2, [DV] −4.75 mm). To identify where *Serpinb2^+^* neurons form synapses, *Serpinb2*-cre mice were unilaterally injected with 0.1-0.15 μl pAAV-hSyn-FLEx-mGFP-2A-Synaptophysin-mRuby in the NAcSh. Then, 10 days after CTB injections and 3 weeks after virus injections, brains tissue was collected and processed for confocal imaging. To aid visualization, images were adjusted for brightness and contrast using ImageJ, but alterations always were applied to the entire image.

### Fiber photometry during feeding

The *Serpinb2^+^, Drd1^+^, Tac2^+^* neuronal dynamics during feeding were measured using fiber photometry. Following injection of an AAV1-hSyn-FLEX-GCaMP7s vector into NAcSh of *Serpinb2*-Cre, *D1*-Cre, *Tac2*-Cre mice, an optical cannula (Ø200 μm core, 0.37 numerical aperture) was implanted 100 μm above the viral injection site. Mice were allowed to recover for 3 weeks and then subjected to behavioral test. GCaMP fluorochrome was excited when GCaMP-expressing neurons were excited, and emission fluorescence was acquired with the RZ10X fiber photometry system, which has built-in LED drivers, LEDs, and photosensors (Tucker-Davis Technologies). The LEDs include 405 nm (as isosbestic control) and 465 nm (for GCaMP excitation). Emitted light was received through the Mini Cube (Doric Lenses) and split into two bands, 420 to 450 nm (autofluorescence) and 500 to 550 nm (GCaMP7 signal). Mice with optical cannula were attached to recording optic cables, and the LED power at the tip of the optical cables was adjusted to the lowest possible (∼20 μW) to minimize bleaching. Mice behaviors were recorded by CCD cameras (Super Circuits, Austin, TX).

For the home cage food consumption recording, fasted mice were habituated to the fiber optic cord for 10 mins in the home cage with normal bedding. After that, we put two pieces of chow (PicoLab Diet, 5053) and one Lego block on the bedding in opposite sides of the cage. During this 10 mins phase, mice can sense the food and freely eat. Behavioral events, such as baseline of free moving, start eating, end eating, sniffing or leaving the Lego block were scored manually and synchronized with fluorescence signal based on recorded videos. The voltage signal data stream was acquired with Synapse software (Tucker-Davis Technologies) and were exported, filtered, and analyzed with custom-written Matlab code. We segmented the data based on individual trials of different events. To calculate ΔF/F, a polynomial linear fitting was applied to isosbestic signal to align it to the GCaMP7 signal, producing a fitted isosbestic signal that was used to normalize the GCaMP7 as follows: ΔF/F = (GCaMP7signal − fitted isosbestic)/fitted isosbestic signal. The Z-score of ΔF/F of heatmap was then calculated as:

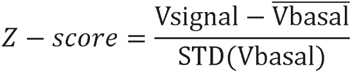

Data is presented using the mean and standard deviation of the signal during the baseline periods (the pooled 10 second time windows before each stimulus). ΔPeak (ΔF/F) was measured by the peak ΔF/F (0s, 10s] minor peak ΔF/F [-10s, 0s]. Time 0s indicates the start time point of each event.

### Behavioral assays

#### Open-field tests (OFT)

A clear box (square 27.3 cm x 27.3 cm square base with 20.3 cm high walls) used for the open field test, and the center zone was 36 % of the total area. Prior to testing, mice were habituated to the test room for at least 20 minutes. Mice were placed in the center of the box at the start of the assay. Movement was recorded using a measurement (Med Associates, St. Albans, VT, ENV-510) 1 hour in 5 mins bins. In addition to regular parameters related to locomotor activity (such as total travel distance, velocity, ambulatory time, resting time), time spent, and distance travelled in the center area of the testing arena were also recorded and analyzed.

#### Conditioned place preference (CPP)

Mice were allowed to freely explore both sides of a custom-made (Med Associates) CPP training apparatus (25 × 19 × 17 cm for L × D × H) for 30 min. Trajectories were tracked by infra-red photobeam detectors, and the travel distance and the duration were recorded to assess their baseline place preference. Mice that showed strong bias (< 25% preference) were excluded from the experiments. Then, for chemogenetic activation or inhibition during CPP formation, these mice were injected with saline (i.p.) and confined to their preferred side of the chamber for 30 min before returned to their home cage. At least four hours later, the same mice received CNO at least 15 min before confined to their non-preferred side of the chamber for 30 minutes. They were then returned to their home cage. The same training with saline and CNO injection were performed for three consecutive days. Twenty-four hours after the final training session, mice were re-exposed to the CPP chamber and allowed to explore both sides of the chamber for 30 min. For the cocaine CPP, mice received CNO at least 15 min before an i.p. injection of 15 mg/Kg cocaine and then were confined to their non-preferred side of the chamber for 30 minutes during conditioning days.

#### Sucrose preference test

Mice were single housed for at least 2Ldays before the tests. For the first 2Ld, animals were habituated with two bottles of water, switching the position of the two bottles at 24Lh. On day 2 night, mice were water deprived for 16Lh and then the test was started on day 3. The test period lasted for 3Lh, during which the mice were exposed to one bottle of pure drinking water and one bottle of drinking water containing 2% sucrose solution (w:v). Bottles were weighed before and after the tests to measure the water consumption. Sucrose preference ratio was calculated by dividing the consumption of sucrose solution by the total consumption of both pure drinking water and sucrose solution.

#### Three-chamber social interaction

Each chamber (30 cm by 30 cm by 30 cm) contains dividing walls with an open middle section to allow for access. Both outer chambers contain wire cups. Mice were given free access to the apparatus for 10 min (absent of other mice) to habituate and confirm initial unbiased preference. The time spent in each chamber was recorded, and the time spent in close interaction with the nose point within 2 cm of the enclosure was also recorded (EthoVision XT 14). To test for sociability, mice were placed into the middle chamber of the apparatus with one outer chamber containing one mouse (stranger 1) confined in wired cup and the other chamber containing a Lego block. For social novelty preference, mice were again placed into middle chamber with one chamber containing the familiar mouse (stranger 1) and the other containing the novel mouse (stranger 2) confined in wired cups. Male familiar and novel mice introduced for assay in social interactions matched the male test subject. For each phase, the test mice explored the entire arena throughout the 10-min trial. The time spent in interacting with the empty wire, stranger 1, and stranger 2 mice during the 10-min session was recorded^69^.

#### Elevated plus maze (EPM)

EPM was used to measure anxiety effect. Before EPM test, mice were brought to the testing room for environmental habituation for at least 30 min. The EPM apparatus is consisted of an elevated platform (80 cm above the floor), with four arms (each arm is 30 cm in length and 5 cm in width), two opposing closed arms with 14 cm walls and two opposing open arms. Mice were individually placed in the center of the EPM apparatus, towards one of the open arms. The mice trajectories were tracked for 5 minutes, and the time spent in the open arms was analyzed using Ethovision XT11 (Noldus).

#### Post-fasted food intake

Mice were individually placed in the home cage and fasted overnight (18 hours) achieving 90% of original body weight. Mice over 92% and lower 88% of original body weight will be excluded in the test. Mice received N-clozapine (CNO, 2mg/mL for hM3Dq group and 5mg/mL for hM4Di group) via i.p. injection and then regular chow pallets (3 g per pellet) were put in the hopper. Three hours later, the remaining food pallets were collected and measured to calculate total amount of food consumed (g). For the leptin titration test, 100 ng, 500 ng, 1 μg of leptin (R&D, Cat# 498-OB) or same volume of saline was delivered through cannula by pump to LH area for 5 mins and waited for another 5 mins before adding pellets. For the leptin treatment test on DREADDs mice, 15 mins after CNO injection, 1 μg or 300 ng of leptin was delivered. For the taCasp3 group, the test was carried out 3 weeks after taCasp3 virus injection.

#### Food place preference

Animals were placed in a custom three-chamber arena (45 × 60 × 35 cm) to assess the time spent in a designated food zone area. The arena contained two 64-cm^2^ food cups in two outer corners of separate chambers. One cup contained standard grain-based chow (Harlan, Indianapolis, IN), while the other cup contained one Lego block. Mice were allowed to explore the arena freely, and spatial locations were tracked using EthoVision XT 10 (Noldus, Leesburg, VA) and CCD cameras (SuperCircuits, Austin, TX). Percentage of food zone time equals time spend in the food zone over total time spend in food zone and non-food zone.

#### Operant behavior

Mice were individually placed in the home cage and kept fasted to maintain the 90% of original body weight. Mice over 92% and lower 88% of original body weight will be excluded in the test. Animals were first given access to 20 mg sweetened chow pellets daily in their home cage before testing, then trained to enter the chamber to retrieve a pellet. Each pellet was delivered 10 s after the prior pellet retrieval. After at least 2 days training and until >30 pellets earned in a single session, animals were trained for the fixed ration 1 (FR1) task, in which each active lever pressing was rewarded with a pellet. A new trial does not begin until animals entered the magazine to retrieve the pellet. Retrieval was followed by a 5 s intertrial interval, after which the levers were reactivated, indicated by a cue light. Training continued until >40 pellets were earned in a single 60 mins session.

#### Progressive ratio

After FR1, FR3, FR5 training sessions, all mice were tested with CNO treatment. For progressive ratio (PR) task, a schedule of reinforcement, each subsequent reward required exponentially more lever pressing based on the increment formula v=(n-1)*c, rounded to the nearest integer, where *n* = number of rewards earned ^70^. 60 mins per session.

#### Optogenetic modulations of post-fasted food intake

Mice received 20 min laser stimulation (4 × 5 min, On-Off-On-Off), and then the remaining food pallets were collected and food intake was measured. For photostimulating ChR2, a 473-nm laser (OEM Lasers/OptoEngine) was used to generate laser pulses (10-15 mW at the tip of the fiber, 5 ms, 20 Hz) controlled by a waveform generator (Keysight), the total duration of the behavioral test was 20 mins, which was divided into four 4 × 5-min epochs (with laser on, off, on and off, respectively). For NpHR photostimulation, a 532-nm laser (OEM Lasers/OptoEngine) generated constant light of 8-10 mW power at each fiber tip.

#### CLAMS recording

Mice were singly housed and habituated to the metabolic cages (CLAMS, Columbus) for at least 3 days before testing under a 12-h light/12-h dark cycle. Mice used in the control and experimental groups (that is, GFP and taCasp3 mice) were age matched. Locomotor activity (infrared beam breaks in the xyz axis), energy expenditure, VO_2_, VCO_2_, RER and food intake were recorded. Data were exported using Clax software v2.2.0 and were analyzed in Prism 9. White and purple represent light (7:00–19:00) and dark (19:00–7:00) cycles, respectively. Energy expenditure, VO_2_ and VCO_2_ data were normalized to body weight. The mice were fed with regular chow (PicoLab rodent diet 20, 5053*; physiological value, 3.43Lkcal g–1). Diets and water were freely available during testing. Gas sensor calibration (CO_2_ and O_2_) of the apparatus was performed before each test. Mouse body weight was recorded before and after every testing session.

### Statistics

All statistical analyses were performed using GraphPad Prism (versions 9) software and fiber photometry results were analyzed by MATLAB.

No statistical methods were used to pre-determine sample sizes. Our sample sizes are similar to those reported in previous publications^71, 72^ ^73^. Mice that, after histological inspection, had the location of the viral injection (reporter protein) or of the optic fiber(s) outside the area of interest, were excluded. Investigators were blinded to the manipulations that subjects had received during recordings and data analysis.

Statistical analyses were two-tailed. Parametric tests, including paired and unpaired *t*-test and one-way ANOVA, were used if distributions passed the Kolmogorov–Smirnov normality test. Normality tests were not performed for one-way ANOVA with missing values. If data were not normally distributed, non-parametric tests were used. One-sample *t*-test was performed to determine whether the group mean differs from a specific value.

For comparisons across more than two groups, one-way ANOVA or repeated-measures one-way ANOVA was performed for normally distributed data, followed by Tukey’s multiple comparison tests; Two-way ANOVA was performed for differences between groups with two independent variables, followed by Sidak’s multiple comparison tests.

All significant *P* valuesL<L0.05 are indicated in the figures. \**P*L<L0.05; \*\**P*L<L0.01; \*\*\**P*L<L0.001; \*\*\*\**P*L<L0.0001. For detailed statistical analysis, see the figure legend with each figure.

## Lead Contact

Further information and requests for resources and reagents should be directed to and will be fulfilled by the Lead Contact, Yi Zhang.

## Data and Code Availability

The MERFISH data are available at the Brain Image Library (https://download.brainimagelibrary.org/fc/4c/fc4c2570c3711952/). Source data, extended data, statements of data and code availability are available at https://doi.org/10.1038/s41593-021-00938-x^30^.For the other custom code that supports the findings from this study are available from the Lead Contact upon request.

## Acknowledgements

We thank Dr. Jeffrey M. Friedman for discussion on some experiments; Dr. Aritra Bhattachejee and Dr.Victoria Honnell for critical reading of the manuscript. We thank the Mouse Behavior Core of Harvard Medical School and its director Dr. Barbara Caldarone for her help. This project was partly supported by 1R01DA042283, 1R01DA050589, and HHMI. Y.Z. is an investigator of the Howard Hughes Medical Institute. This article is subject to HHMI’s Open Access to Publications policy. HHMI lab heads have previously granted a nonexclusive CC BY 4.0 license to the public and a sublicensable license to HHMI in their research articles. Pursuant to those licenses, the author-accepted manuscript of this article can be made freely available under a CC BY 4.0 license immediately upon publication.

## Contributions

Y.Z. conceived the project; Y.L., and Y.Z. designed the experiments; Y.L. performed most of the experiments. Y.W. and Z.-D.Z. helped with the fiber photometry and data analysis. G.X. helped with the catheter administration, CLAMS recording and staining. C.Z. analyzed MERFISH data. R.C. initiated the Serpinb2-Cre mouse generation. Y.L., Y.W., Z.-D.Z., G.X. and Y.Z. interpreted the data; Y.L. and Y.Z. wrote the manuscript with input from Z.-D.Z and R. C.

## Ethics declarations

## Extended Data Figures

**Extended Data Fig. 1.**
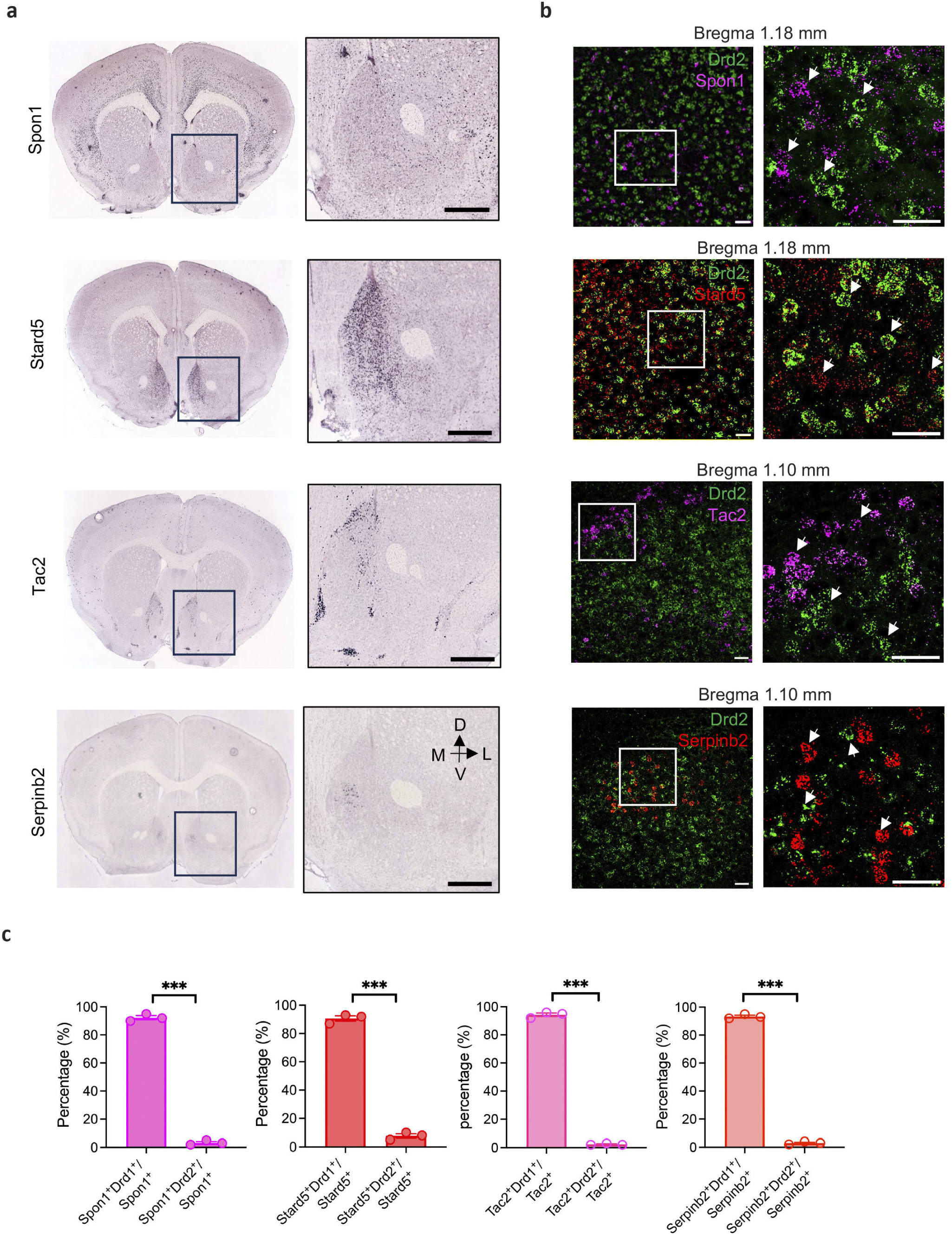
The D1 subtype neuron marks do not overlap with Drd2 in MSN NAcSh. **a,** ISH images showing the spatial distribution of *Spon1*, *Stard5*, *Tac2* and *Serpinb2* markers of certain D1 MSN subtypes, in mouse NAc. Boxed regions in left panels are enlarged and shown in the right panels. The data are from the Allen Mouse Brain Atlas. Scale bars, 500Lµm. **b,** smFISH confirms the expression of MSN D1 subtype-specific markers in the NAcSh do not overlap with Drd2. Boxed regions in left panels are enlarged and shown in the right panels. Three independent experiments were performed with similar results. Scale bar, 50 µm. **c,** The percentage of NAc^Spon1^, NAc^Stard5,^ NAc^Tac2^ and NAc^Serpinb2^ cells overlapping with *Drd1*^+^ or *Drd2*^+^ cells in the NAcSh. For **b**–**c**, every fourth of 16-µm brain sections was counted. *n*L=L3 mice for all groups. All statistical tests are Paired two-tailed *t*-tests. Data are mean ± SEM. \*\*\**P*L<L0.001.

**Extended Data Fig. 2.**
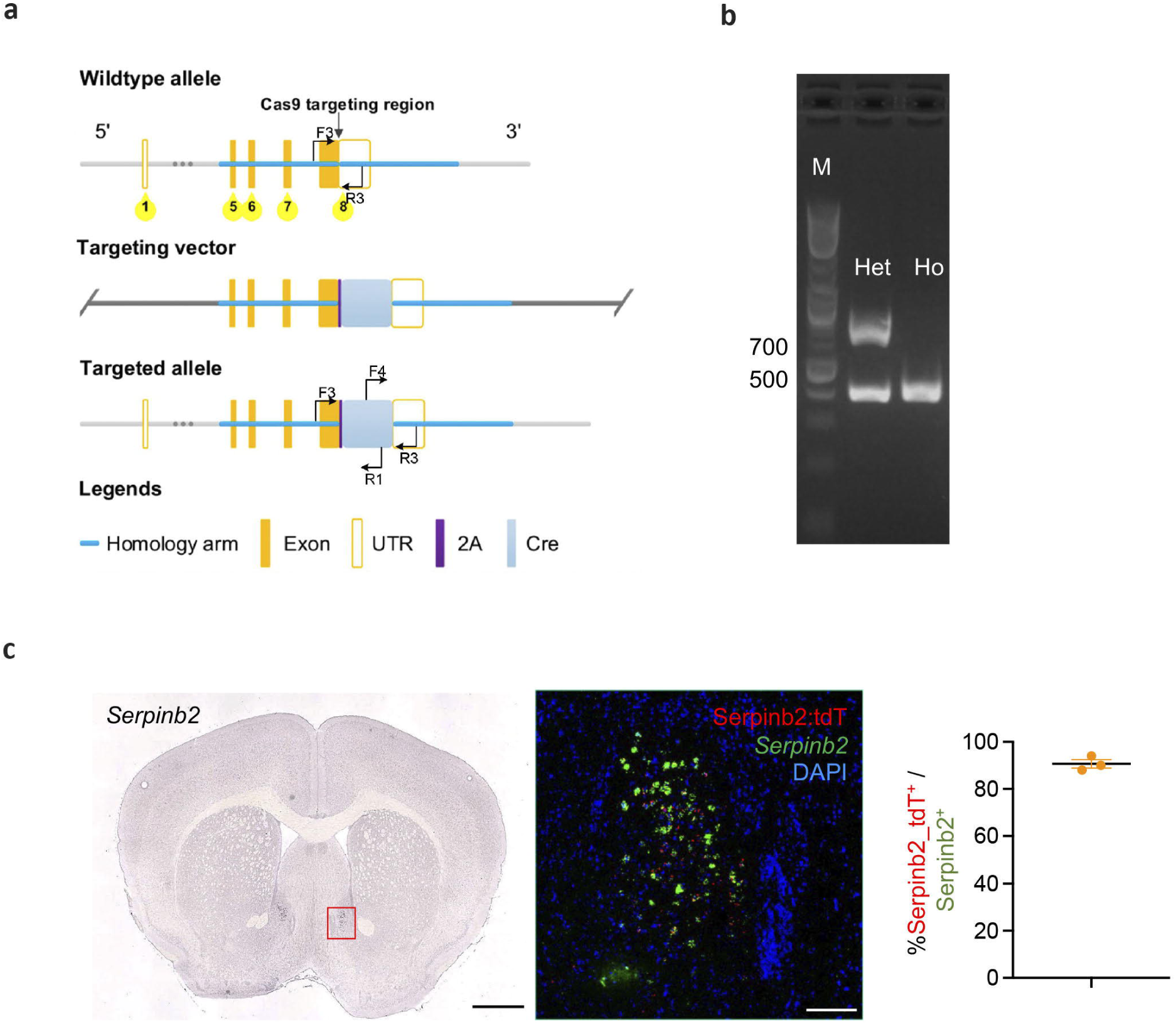
Generation and validation of a *Serpinb2*-Cre mouse line. **a,** Diagrams showing the gene targeting strategy. **b,** Genotyping by PCR. Homozygotes: 413 bp. Heterozygotes: 413 bp/772 bp. **c,** Left, in situ hybridization (ISH) data of *Serpinb2* from Allen Brain Atlas. Scale bar: 1000 μm. Middle, co-localization of endogenous *Serpinb2* (green), tdTomato (red) and DAPI (blue). Scale bar: 100 μm. Right, quantification of tdTomato^+^ neurons among all *Serpinb2*^+^ neurons. *n*L=L3 mice

**Extended Data Fig. 3.**
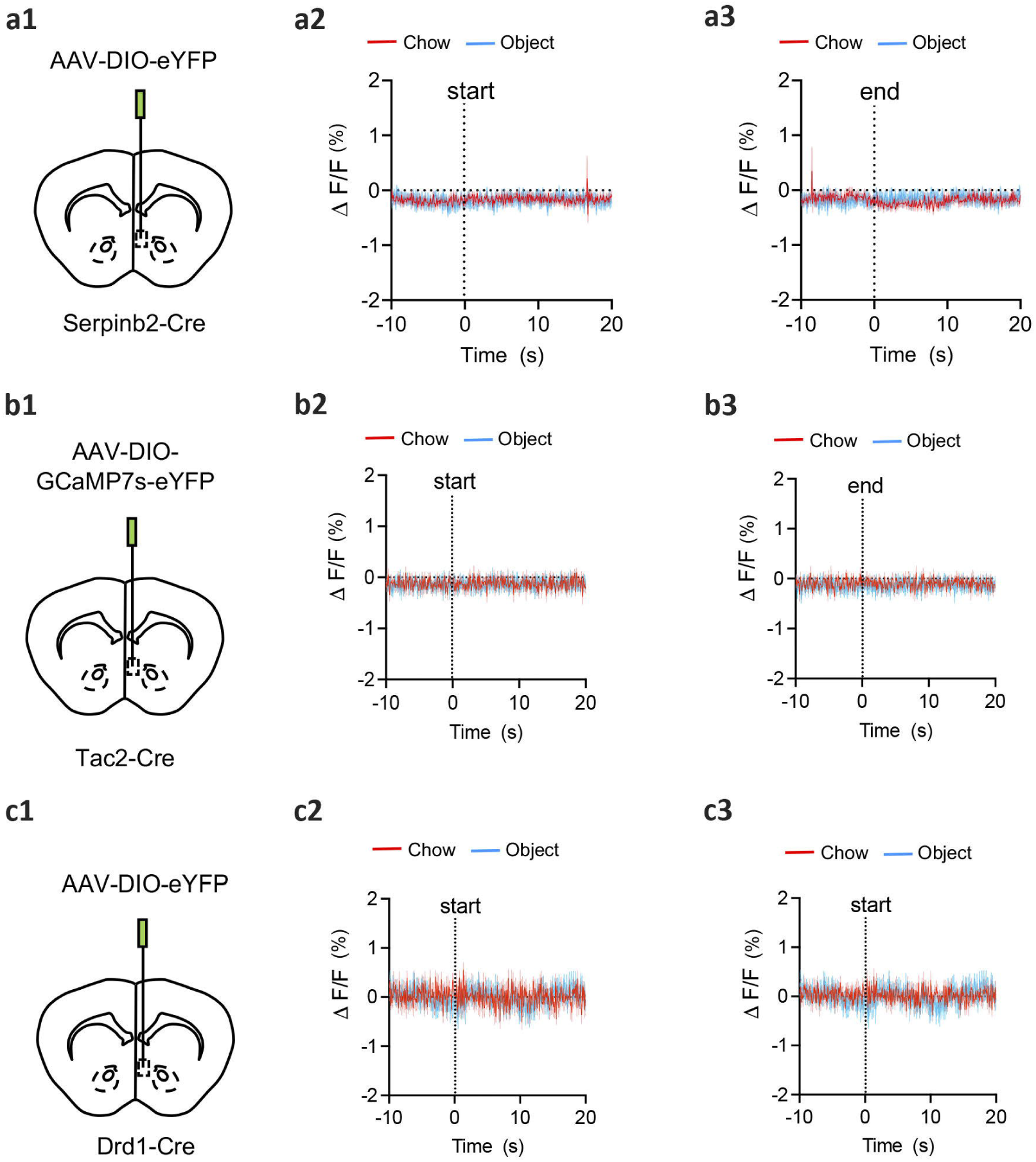
*Tac2*^+^ neurons do not respond to feeding behavior. **a1-a3,** The injection site of the AAV-DIO-eYFP virus in NAcSh of *Serpinb2*-Cre mice as negative control (**a1**). Average peri-stimulus histograms of Ca^2+^ signals of *Serpinb2*^+^ neurons. Dash line, eating start point (**a2**). Average peri-stimulus histograms of Ca^2+^ signals of *Serpinb2*^+^ neurons. Dash line, eating end point (**a3**). n=5 mice. **b1-b3,** similar as **a1-a3** but with AAV-DIO-GCaMP7s virus into *Tac2*-Cre mice NAcSh. The Ca^2+^ signal recording is the same as in **a2-a3**. n=5 mice. **c1-c3,** similar as **a1-a3** but in *Drd1*-Cre mice. N=4 mice. Data are represented as mean ± SEM.

**Extended Data Fig. 4.**
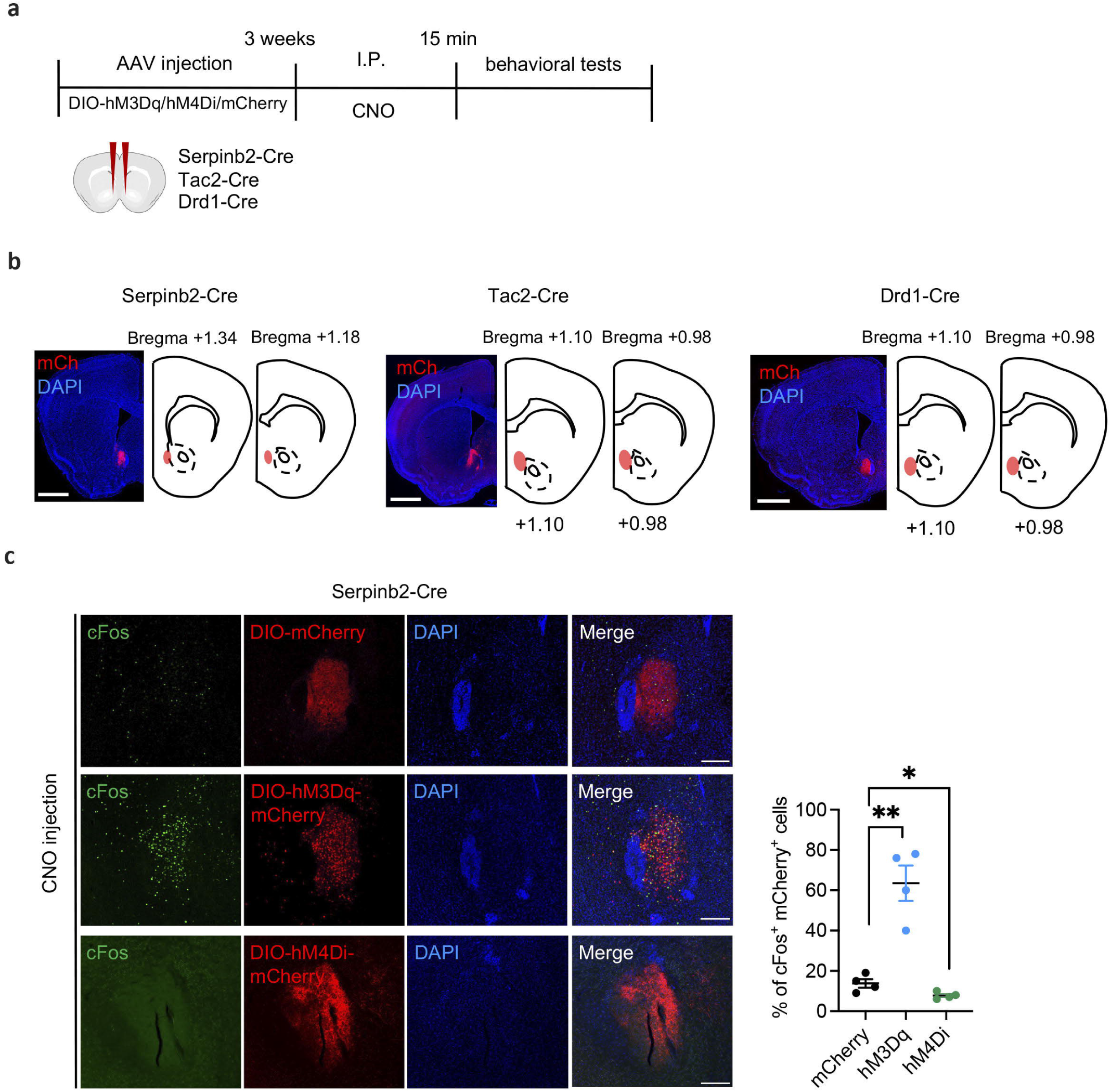
Validation of chemogenetic manipulation. **a,** Experimental scheme of chemogenetic manipulation. **b,** Validation of virus expression in *Serpinb2*-Cre, *Tac2*-Cre and *Drd1*-Cre mice. Scale bar: 500 μm. **c,** *cFos* induction after intraperitoneal injection of ligand CNO in mCherry-expressing, hM3Dq-mCherry-expressing and hM4Di-mCherry-expressing mice. The ratio of cFos^+^/mCherry^+^ cells in all mCherry^+^ cells was calculated and shown on the right panel. Scale bar: 100 μm. **, p<0.01, *, p<0.05; ns, P > 0.05, one-way ANOVA test. Data are represented as mean ± SEM.

**Extended Data Fig. 5.**
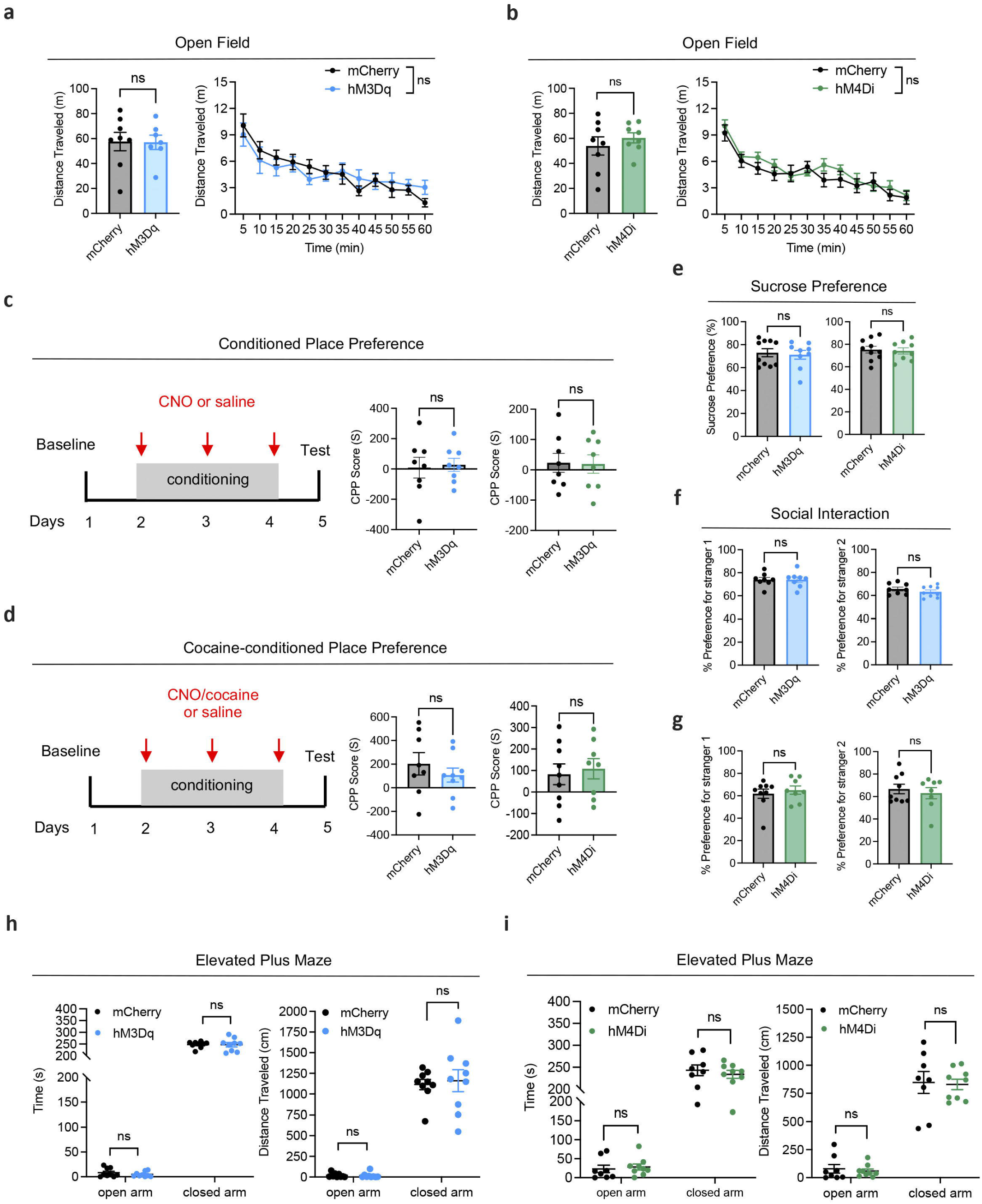
*Serpinb2*^+^ neuronal activity does not affect locomotion, drug seeking, social, anhedonic or anxiety behavior. **a, b,** Open field test for the effect of *Sepinb2*^+^ neuronal activation (hM3Dq) (a) or inhibition (hM4Di) (b) on the total distance traveled in the 1-hour post-treatment period after chemogenetic manipulation of *Sepinb2*^+^ neurons (left) or the distance traveled in 5-min time bin (right). ns, p>0.05, left, unpaired t-test; right, two-way ANOVA. **c,** Left, illustration of the two-chamber CPP paradigm. Right, CPP with chemogenetic activation (hM3Dq) or inhibition (hM4Di) of *Sepinb2*^+^ neurons. CPP scores were calculated by subtracting the time spent in the preconditioning phase from the time spent in the postconditioning phase. ns, p>0.05, unpaired t-test. **d,** The same as in panel **c** except conditioned with cocaine. **e,** Sucrose preference test for the effect of *Sepinb2*^+^ neuronal activation (hM3Dq) (left) or inhibition (hM4Di) (right). ns, p>0.05, unpaired t-test. **f, g,** 3 chamber social interaction test for the effect of *Sepinb2*^+^ neuronal activation (hM3Dq) (up) or inhibition (hM4Di) (bottom). ns, p>0.05, unpaired t-test. **h, i,** Elevated plus maze test for the effect of *Sepinb2*^+^ neuronal activation (hM3Dq) (h) or inhibition (hM4Di) (i) on the time spent (left) or distance traveled (right) in open arm and closed arm of the 5-min post-treatment period after chemogenetic manipulation of *Sepinb2*^+^ neurons. ns, p>0.05, unpaired t-test. Data are represented as mean ± SEM.

**Extended Data Fig. 6.**
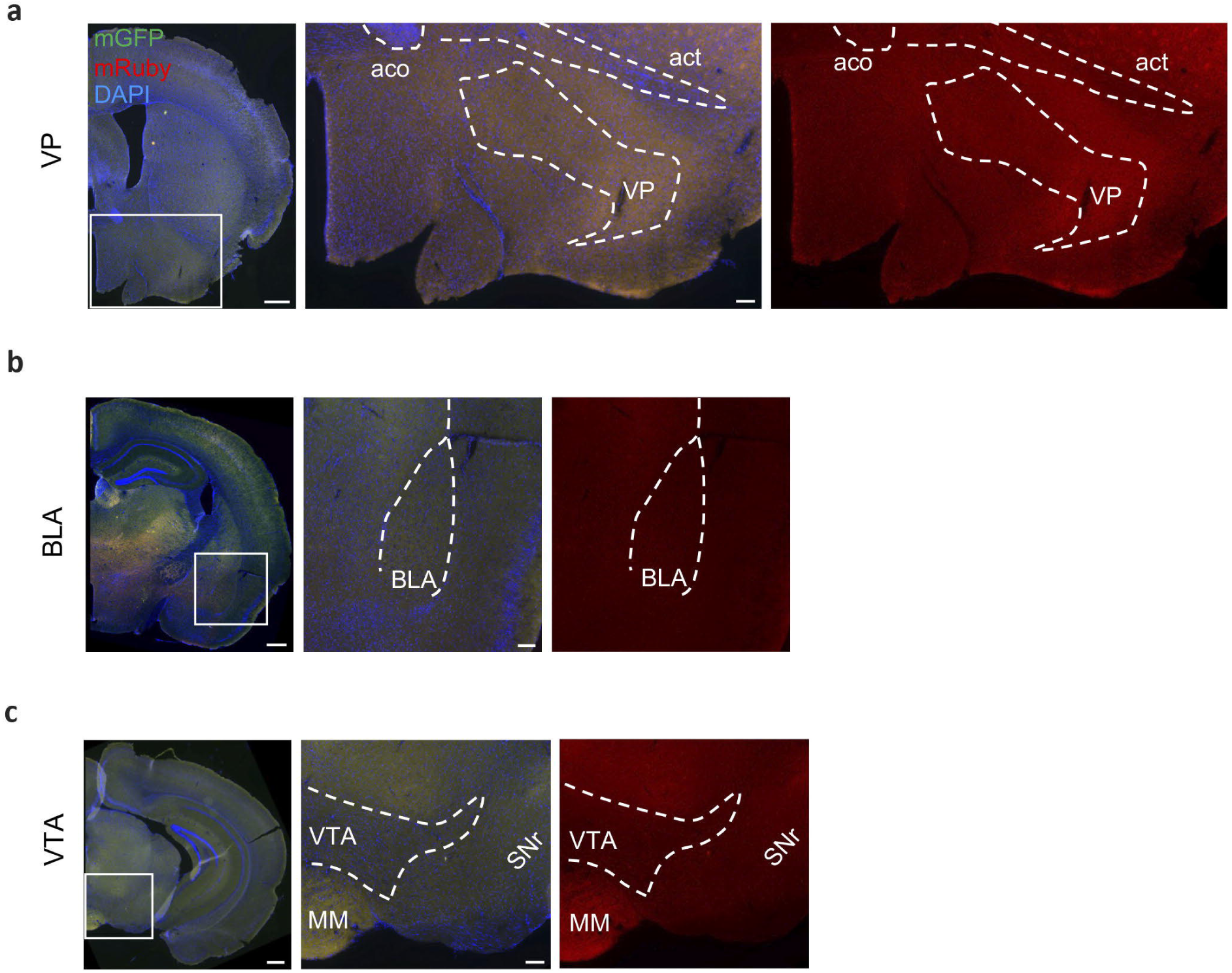
Projection mapping of *Serpinb2*^+^ neurons. **a-c,** No detectable signals of mGFP or mRuby in VP (**a**), BLA (**b**) or VTA regions (**c**). aco, anterior commissure; act, anterior commissure, temporal limb; BLA: Basolateral amygdala; MM, medial mammillary nucleus; SNr, substantia nigra, reticular part. Scale bars, left, 500 μm; others, 100 μm.

**Extended Data Fig. 7.**
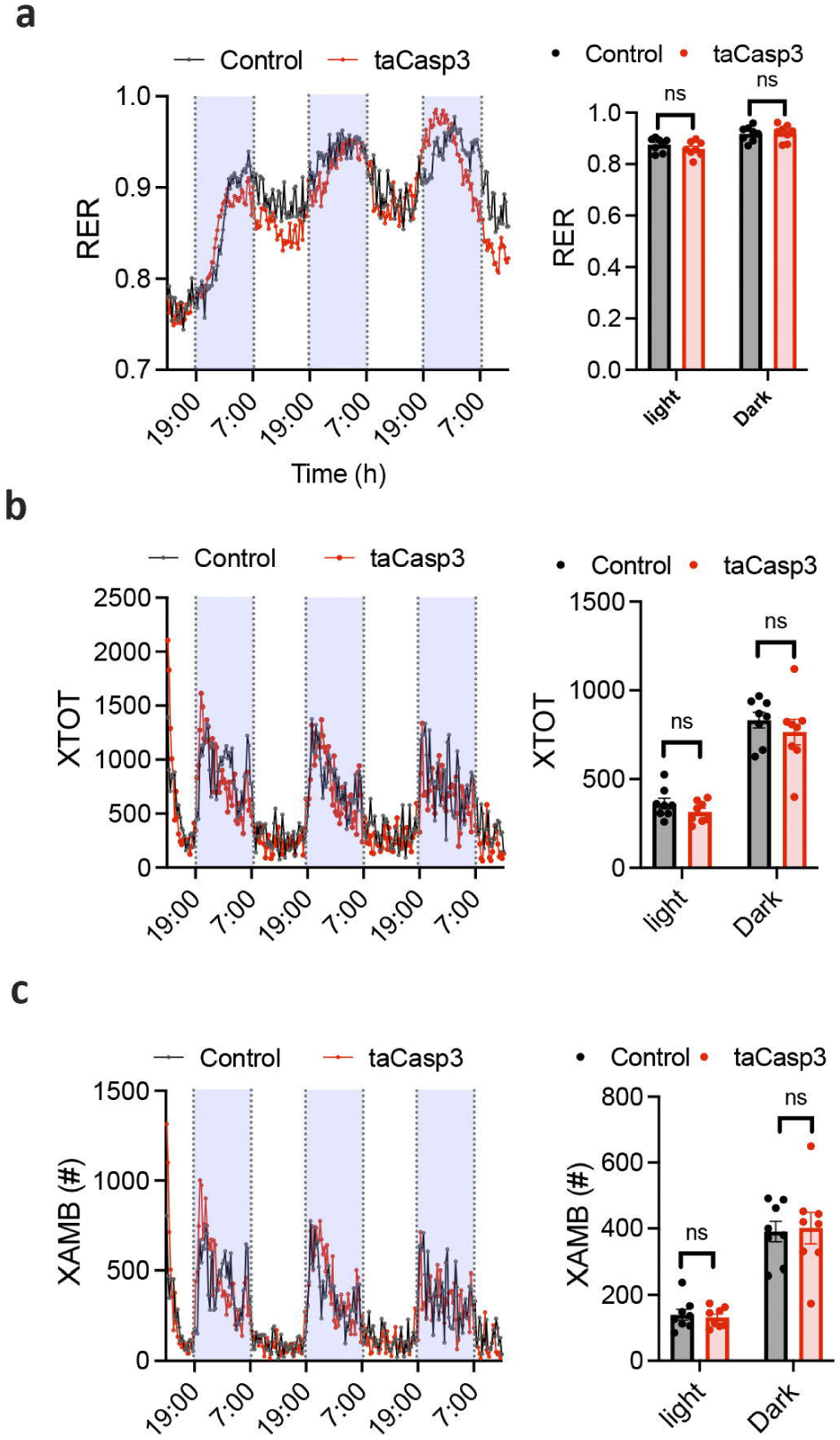
Ablation of *Serpinb2*^+^ neurons does not affect RER and locomotion. **a,** Respiratory exchange ratio (RER) of mice GFP-expressing control (n = 8) and taCasp3 (n = 8) mice fed chow. **b,** total activity, GFP-expressing control (n = 8) and taCasp3 (n = 8) **c,** ambulatory movement, GFP-expressing control (n = 8) and taCasp3 (n = 8). Data are presented as means ± SEM. ns, p>0.05, unpaired t-test.

